# Pooled CRISPRi screening reveals fungal-specific vulnerabilities across environments and genetic backgrounds

**DOI:** 10.64898/2025.12.19.695556

**Authors:** Lauren F Wensing, Philippe C Després, Desiree Francis, Meea Fogal, Anthony Hendriks, Nicholas C Gervais, Clara Fikry, Abdul-Rahman Adamu Bukari, Aleeza C Gerstein, Christina A Cuomo, Rebecca S Shapiro

## Abstract

The rising rate of drug-resistant fungal infections and the emergence of fungal pathogens with intrinsic resistance phenotypes are a growing concern. The close evolutionary distance between mammals and fungi complicates the design of new antifungals and increases the chances of toxic off-target effects. As such, antifungal drug development usually focuses on fungal-specific proteins when considering potential new targets. Ideal drug targets should mediate essential cell processes and be highly sensitive to inhibition. Targeted gene repression can serve as a model for drug-mediated inhibition and for determining the dosage-sensitivity profile of genes of interest. In the fungal pathogen *Candida albicans*, classical approaches for gene repression can be labour-intensive and limited to one genetic background due to low throughput. Here, we adapt pooled CRISPRi screening in *C. albicans* for the first time and exploit this technique for large-scale functional genomic analysis. Through pooled CRISPRi screening, we test the repression sensitivity of over a hundred essential genes conserved in fungi but absent in humans, and successfully identify highly dosage-sensitive genes across multiple cell components and pathways. By extending our analysis to ten diverse environmental conditions, we show how the environment influences dosage-sensitivity profiles. Finally, we extend our experiments to two clinical drug-resistant *C. albicans* strain backgrounds and demonstrate that many of the fitness defects we observed are conserved in resistant clinical isolates. Together, our results highlight a set of genes that are highly dosage-sensitive across different genetic and environmental contexts, making them attractive targets for further investigation. By facilitating rapid, efficient large-scale functional genomics assays across diverse genetic backgrounds, CRISPRi pooled screening will open new frontiers in *C. albicans* biology.

## Introduction

Over recent years, fungal infections have become more prevalent due to environmental changes and an increase in the immunocompromised population, a trend expected to continue in the coming decades^1^. One of the most medically relevant human fungal pathogens is *Candida albicans*, which is the fourth most common cause of hospital-acquired bloodstream infections and results in roughly a million deaths each year^2^. Current treatment options for invasive *Candida* infections are limited to three classes of antifungal drugs, with two of these, azoles and echinocandins, being dominant in therapeutic management. While these therapeutics have been broadly effective, the increased occurrence of antifungal resistance, as well as cases of cross-resistance^3^, has underscored the urgent need for new antifungal drugs. The close evolutionary relationship between fungi and humans has significantly complicated this process. For example, promising molecules with antifungal effects, such as rapamycin (a TOR complex inhibitor) and FK506 (a calcineurin inhibitor), were later found to have potent immunosuppressive effects in mammals due to the conservation of both complexes, precluding their use to treat fungal infections^4^. To avoid crosstalk with human cell components, fungal-specific cell components must therefore be prioritized as targets for future antifungal development.

In the genomes of microbial pathogens, essential genes have commonly been the focus of antimicrobial development efforts as they, by definition, directly affect the cell viability and reproduction^5^. However, knowing that a given gene is essential provides no information about what fraction of its activity must be inhibited to significantly impede growth (Figure 1a). This relationship between gene dosage and fitness, often referred to as gene dosage-sensitivity, in turn mediates the dose-response profile to an inhibitor. The fraction of gene activity that must be reduced by mutations or inhibited by drugs to cause a fitness defect can be influenced by expression levels above the basic requirements for optimal fitness^6^ and by downstream buffering by metabolic networks^7^. Dosage sensitivity can vary immensely between genes^8^ and has not been characterized for most of the *C. albicans* genome. A gene for which only a slight reduction in activity translates to reduced fitness should be more sensitive to drug inhibition. For example, the dosage of haploinsufficient genes has low buffering capacity, so that reducing the effective concentration of the gene product by deleting one out of two alleles results in fitness defects or drug sensitization^9^. Importantly, dosage-sensitivity is not captured by *in vitro* enzymatic assays alone, which isolate the protein of interest outside of its cellular context. Finding and prioritizing highly dose-sensitive targets enables the use of lower drug doses, improving therapeutic safety and allowing for less stringent optimization of compounds during drug development. The current lack of information on dosage-sensitivity profiles for fungal pathogens’ genes limits our ability to achieve this. Importantly, successful antifungal targets must remain effective across a range of environmental conditions and genetic backgrounds, including drug-resistant isolates, as these contexts can profoundly influence gene-fitness relationships^10^. However, for essential genes, filling this knowledge gap is complicated by the inherent challenges of manipulating them genetically.

**Figure 1:**
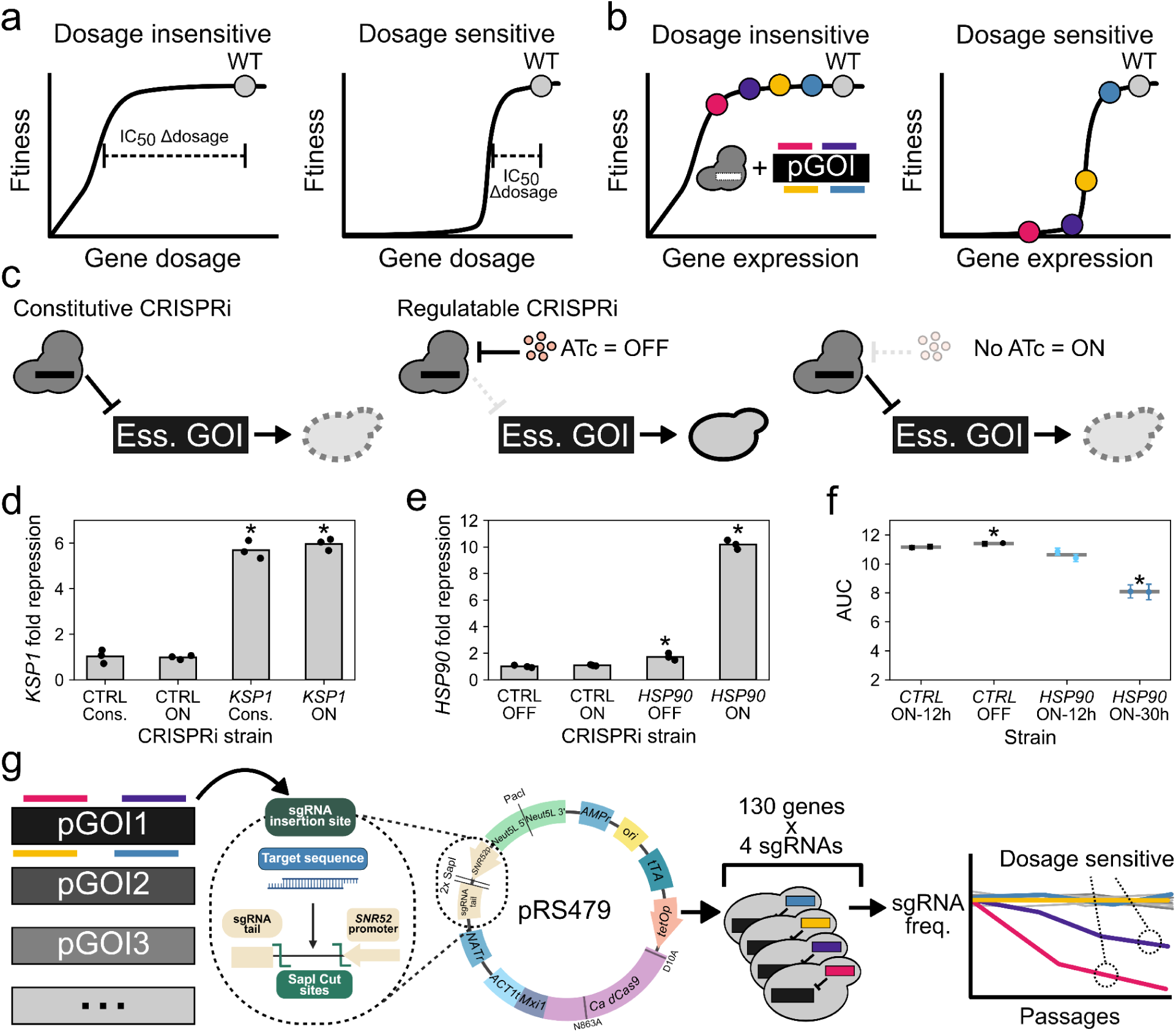
Regulatable CRISPRi for pooled screening in *Candida albicans*. **a)** The dosage sensitivity profile of a gene of interest (GOI) impacts druggability. For dosage-insensitive genes, even a large reduction in the effective concentration of gene products can result in only minor fitness effects, while only a slight reduction in the abundance of a dosage-sensitive product leads to large fitness effects. **b)** Gene knockdown with CRISPRi reduces expression levels by targeting a transcriptional repressor to the promoter of a gene of interest (pGOI), with each different sgRNA potentially leading to different levels of depletion and mimicking drug repression. As such, sgRNAs targeting the same gene at different degrees of repression can identify repression-sensitive genes. **c)** The fitness of CRISPRi strains with sgRNAs targeting essential GOIs is immediately impacted if dCas9-Mxi1 expression is constitutive, leading to difficulties and premature dropouts when constructing strains. Placing *dCas9-Mxi1* under the control of a *tetO* promoter/repressor system prevents immediate fitness effects, which occur only when switching cells to media without ATc, thereby lifting repression of the CRISPRi system. **d)** CRISPRi repression of *KSP1* in strains transformed with either the constitutive (cons.) or the regulatable system in its active state (ON) with the same sgRNA. Stars represent significant differences between the *KSP1* targeting strains and the control strains (Cons.: p = 0.0001, ON: p = 0.0002, Welch’s t-test). **e)** CRISPRi repression of *HSP90* by the regulatable system when it is inactive (OFF) or active (ON). Stars represent significant differences with the OFF control strain (*HSP90* OFF: p = 0.03, *HSP90* ON: p = 0.0002, Welch’s t-test) **f)** Growth effect of *HSP90* repression using the inducible system after different ATc washout periods. Stars represent significant differences with the WT-12h control strain (CTRL OFF: p = 0.019, *HSP90* ON-30h: p = 0.00026, Welch’s t-test). **g)** Workflow for pooled CRISPRi screening of fungal-specific essential genes in *C. albicans*. For each gene of interest, four sgRNAs were designed and cloned into the regulatable CRISPRi integration vector from an oligonucleotide pooled library. This plasmid library was then batch transformed into fungal cells under ATc repression. The resulting library can then be used for pooled screening, in which changes in relative sgRNA abundance measured by high-throughput sequencing are used to infer fitness effects and repression sensitivity.

The advent of CRISPR technologies in *Candida* species presents new opportunities for a deeper characterization of essential genes and their function beyond direct editing^11–13^. CRISPR interference (CRISPRi), in which a transcriptional repressor is fused with a catalytically dead Cas9 (dCas9) enzyme, is targeted to a promoter by a sgRNA and can be used to knock down gene expression of genes of interest in a programmable manner^14,15^. This strategy provides an alternative to locus-specific promoter replacement, enabling tunable gene repression while simplifying strain construction and preserving the native genomic context^16–18^. In addition, CRISPRi systems allow the design of unique sgRNAs targeting the same promoter, each of which induces a different level of repression depending on the guide and promoter sequence properties. Accordingly, genes that exhibit fitness defects when repressed by multiple distinct sgRNAs can be interpreted as repression-sensitive, based on their vulnerability to a broad range of decreases in expression, mimicking drug inhibition and suggesting dosage-sensitivity (Figure 1b). Another advantage of CRISPR approaches is their high scalability. High-throughput CRISPRi libraries have been used in a diverse range of eukaryotic organisms to identify key cellular components through pooled competition assays^19,20^, but this approach has yet to be extended to fungal pathogens. In these experiments, high-throughput sequencing of the sgRNA locus can be used to infer the fitness effects of thousands of perturbations in a single test tube. In *C. albicans*, such approaches could enable rapid characterization of genes of interest and are well-suited to the study of essential cell components.

Here, we develop and validate a first-of-its-kind regulatable system for pooled CRISPRi screening in *C. albicans*. Using this platform, we interrogate the repression sensitivity of 130 putative essential genes that are conserved in fungi but absent in humans, both in standard laboratory conditions and across ten diverse environmental contexts. We further test the impact of genetic background on repression-sensitivity profiles by repeating our screen with two clinical antifungal-resistant isolates. We find that our pooled CRISPRi approach can robustly detect genes that are sensitive to dosage perturbations. We identify a group of 16 genes that showed significant repression sensitivity under all experimental and genetic contexts, nominating them as high-priority candidates for future antifungal drug development. More broadly, our approach provides a rapid, scalable, and portable method for creating large knockdown mutant libraries that can be readily applied to elucidate gene function under different environmental conditions and across different genetic backgrounds in *C. albicans*.

## Results

### Design and construction of the targeted essential gene library

To enable scalable, pooled repression of essential genes, we built on our previously established CRISPR interference (CRISPRi) framework in *C. albicans*, in which a catalytically dead Cas9 (dCas9) fused to the transcriptional repressor Mxi1 is directed to gene promoters by a single guide RNA (sgRNA). In this system, all CRISPRi components are encoded on a single integrating plasmid, simplifying strain construction and enabling pooled library generation^14^. In our previously published *C. albicans* CRISPRi system, dCas9–Mxi1 was expressed constitutively following integration at a safe-harbour locus. However, controlling the timing of gene repression is critical when interrogating essential genes, particularly to avoid premature depletion of strains before the start of competitive growth assays (Figure 1c). Thus, to create a regulatable gene repression system, we redesigned our vector to include a tetracycline (tet)-regulated transcription factor and an associated promoter (*tetO*) controlling dCas9 expression^16^. This *tetO* promoter is a *C. albicans*-optimized tet-off system in which the addition of tetracycline or an analog, such as anhydrotetracycline (ATc), represses dCas9 expression. In the absence of ATc, dCas9 is expressed, and the gene targeted by the sgRNA is repressed. We validated this new system using one non-essential gene, *KSP1* and one essential gene, *HSP90*. The regulatable expression system achieved comparable repression efficiency compared to constitutive expression (Figure 1d). In addition, the promoter exhibited low expression leakage, with minimal repression of *HSP90* when turned off and a strong effect when turned on (Figure 1e). Knockdown of *HSP90* only led to growth effects when ATc was absent over multiple generations, with minimal expression costs (Figure 1f). The regulatable system thus provides control over the timing of repression-induced effects, allowing for better decoupling of strain construction and fitness measurements. This modified CRISPRi system, which is well-suited to study essential genes, now enables us to scale up and develop an efficient strategy to generate a pooled *C. albicans* mutant library, supporting large-scale, targeted forward genetic screening of essential genes (Figure 1g).

We next sought to use our regulatable CRISPRi system to identify repression-sensitive essential genes in *C. albicans* at high-throughput. We focused on a list of 130 putatively essential genes identified in a transposon mutagenesis screen (TN-Seq) of a haploid *C. albicans* model (Table S1). In addition to being depleted for transposon insertions, these genes were conserved across three evolutionarily distant fungal pathogens (*Aspergillus fumigatus*, *Histoplasma gondii*, and *Cryptococcus neoformans*) and lacked close homologs in humans^21^. We used the sgRNA design principles outlined in a previous CRISPRi screen in *Saccharomyces cerevisiae*^22^ to select sgRNA target sequences. Briefly, we selected four guides targeting the promoter of each gene, prioritizing sgRNAs with higher predicted EuPaGDT efficiency scores^23^. We also included 30 randomized sgRNAs sequences as negative controls (control sgRNAs). The library was synthesized as a pool of single-stranded oligonucleotides, which was then amplified and cloned *en masse* into the CRISPRi plasmid. We transformed the plasmid pool into *C. albicans* laboratory strain SC5314. For both bacterial and fungal library generation steps, we validated that library sequencing recovered the vast majority of designed sgRNAs, with ∼85% of the designed sgRNAs being present in the fungal pooled library (Figure S1). Our optimized pipeline for pooled strain construction enables the generation of any screening-ready *C. albicans* CRISPRi library in as little as seven days (Supplementary Note 1). We have made the regulatable CRISPRi vector publicly available on Addgene (pRS479, #250493), thus providing a new, cost-effective and rapid option for high-throughput functional genomics in *C. albicans* research.

### High-throughput screening of essential genes in standard laboratory conditions

Having built our CRISPRi strain library, we next established a pipeline to screen our mutant library for growth defects under standard laboratory conditions. We grew the library in rich media (YPD) with or without ATc over three 12-hour passages, spanning ∼21 generations, and tracked changes in the relative abundance of mutant strains compared to the starting inoculum population using high-throughput sequencing of the sgRNA locus. Log_2_ fold-change (Log_2_FC) values were well correlated between replicates (ρ > 0.82, p < 0.0001 for all correlations), showing that the pooled competition assay generated reproducible outputs (Figure S2). When CRISPRi repression was turned on (CRISPRi-ON), the relative abundance of some sgRNAs targeting essential gene promoters decreased sharply compared to the randomized controls, while abundances remained mostly unchanged when the system was kept inactive (CRISPRi-OFF) (Figure 2a, Figure S3). Using the Log_2_FC values of the random sgRNAs as a control distribution, we found 96 sgRNAs were significantly depleted in abundance at the final time point, which we refer to as hits (Figure 2b, Figure S4). To validate that negative Log_2_FCs were indeed associated with reduced fitness, we constructed 10 strains with sgRNAs targeting *MRLP33*, *MMM1,* or *ASK1*, candidate genes from this screen with three or more hit guides. Both *MRPL33* and *MMM1* are linked with mitochondrial function, while *ASK1* is a subunit of the DASH complex involved in chromosome segregation. In most cases (9/10), screen hit sgRNAs induced a measurable growth defect in smaller-scale growth curve assays (Figure 2c), validating the phenotypes identified in the large-scale assay. This demonstrates the ability of our novel, pooled CRISPRi screening assay for rapid and scalable analysis of *C. albicans* strains libraries.

**Figure 2:**
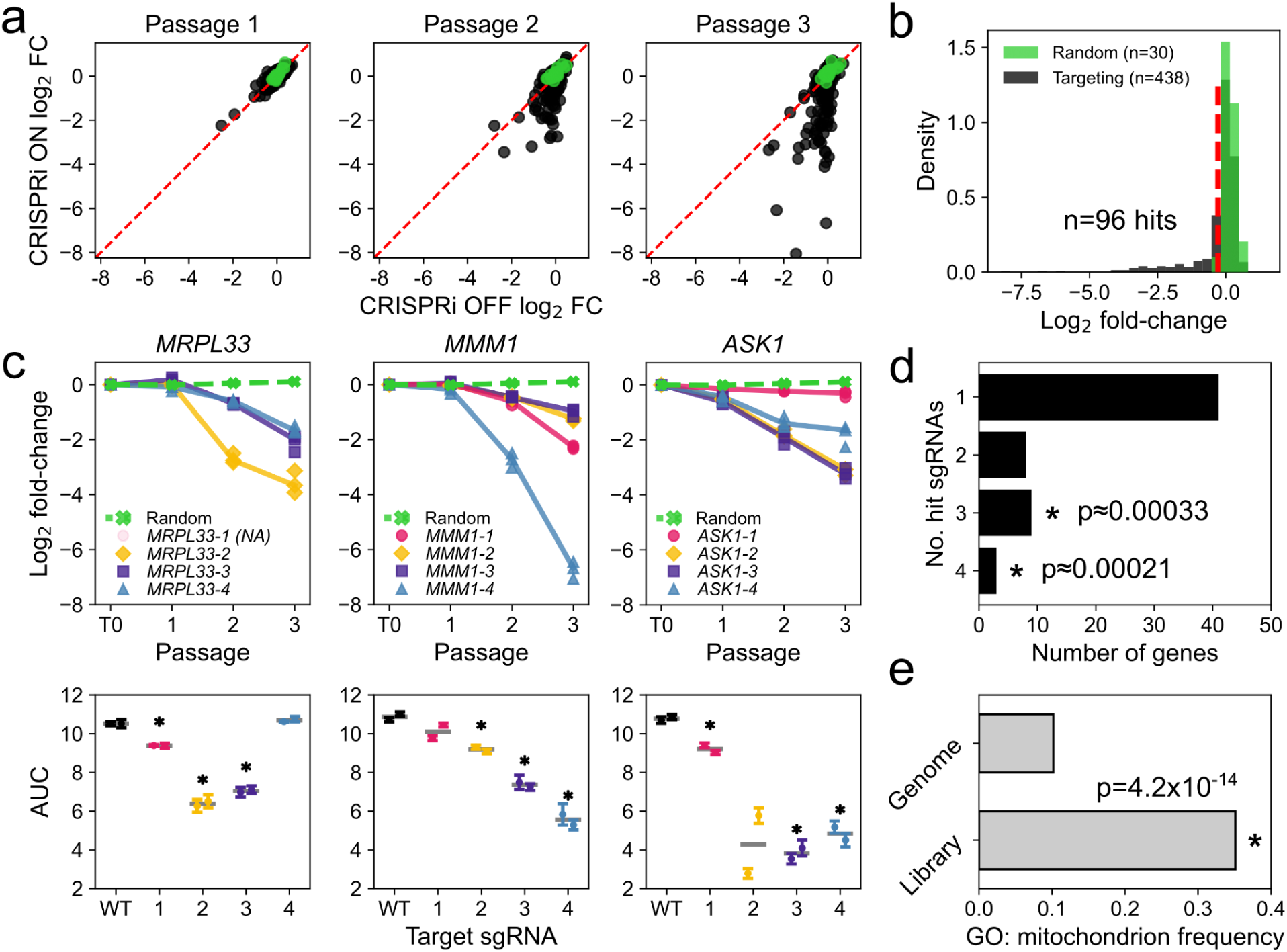
Pooled CRISPRi screening of essential genes in *C. albicans* identifies repression sensitivity phenotypes. **a)** Changes in relative abundance of sgRNAs in the library throughout the timecourse assay, with (ON) or without (OFF) CRISPRi repression. Randomized sgRNAs (n=30) are shown in green, and essential gene targeting sgRNAs are shown in black. The red dotted line shows a hypothetical 1:1 relationship. **b)** Abundance change distribution at the final time point for the CRISPRi-ON condition. The red line indicates the significance threshold corresponding to FDR=5%. **c)** Abundance changes of sgRNAs targeting *MRPL33*, *MMM1* and *ASK1* in the timecourse assay, with the corresponding AUC measurement obtained in growth curve assays. The grey bars show the mean of two biological replicates, which are shown overlaid with error bars corresponding to the 95% confidence interval of 5 technical replicates. Stars indicate strains with a significant change in growth (p<0.05, Welch’s t-test). **d)** Number of hit sgRNAs per gene in the timecourse assay, for genes with at least one hit. Empirical p-values were obtained by sampling 96 hits over the 438 sgRNAs measured over 100 000 permutations to estimate the expected frequency of genes with 3 or 4 hits. **e)** Frequency of mitochondrial localization annotation (GO:0005739) across the *C. albicans* genome (n=6473) and within the fungal-specific gene set (n=131). The difference in annotation frequency in both categories was tested using Fisher’s exact test.

Amongst mutant strains with growth defects, we found additional instances in which three or four of their four designed sgRNAs were significantly depleted, yielding a total of twelve enriched genes. A higher fraction of sgRNAs that result in fitness defects is unlikely to occur by chance, suggesting greater repression sensitivity for these targets (Figure 2d). These twelve genes include five that are predicted by *S. cerevisiae* orthology to encode part of the mitochondrial ribosome: *MRPL33*, *C1_12610W* (S.c.: *MRPL15*), *C2_07680W* (S.c.: *MRPL7*), *CR_01370C* (S.c.: *MRPS28*), and *CR_04140W* (S.c.: *NAM9*), of which four remain uncharacterized in *C. albicans*. These five proteins do not cluster spatially within the S. cerevisiae mitoribosomal structure (Figure S5) but might represent different, potentially rate-limiting components of complex assembly. In *S. cerevisiae*, *MRPL33*, *MRPL15*, and *MRPL7* are essential components of the large subunit of the mitoribosome that join the complex in the early to intermediate stages of assembly^24^. Two other highly repression-sensitive genes, *MMM1* and *C1_00510W* (S.c.: *ECM31*), are also predicted to be localized to the mitochondria^25^. This reflects a high enrichment of the initial gene set for mitochondrial genes and mitoribosomal components (Figure 2e). Collectively, these observations suggest that mitochondrial pathways, particularly components of the mitoribosome, are among the most repression-sensitive components in our screen and may therefore constitute especially promising targets for antifungal inhibition.

### Screening in multiple environments identifies a subset of core repression sensitive genes

Our initial pooled-library experiment enabled us to identify repression sensitive essential genes under standard laboratory conditions. However, environmental factors can affect the fitness contribution of genes, potentially leading to condition-dependent essentiality^10,26^. Ideally, candidate antifungal targets should show strong repression sensitivity across all environments. To test this, we screened our CRISPRi library across 11 alternative environments, spanning different nutrient sources and stresses, and monitored changes in the relative abundance of mutant strains via sequencing (Figure 3a). We found that changes in mutant strain abundance were highly correlated within conditions (within condition inter-replicate median ρ = 0.80-0.96, p < 0.0001 for all correlations; Figure S6), supporting the high reproducibility of the approach. Different signal-to-noise ratios across screening conditions resulted in variation in the number of hits identified in each screen, prompting us to exclude the tunicamycin condition from downstream analysis (Figure 3b). Mapping CRISPRi-induced defects across conditions, we found that most phenotypes were broadly identical across conditions (e.g. *MMM1*, *ASK1*, *C2_07680W*, *C2_03560C*, Figure S7), with a few cases of condition-dependent fitness defects (Figure 3c). Overall, these results demonstrate the applicability of our pooled CRISPRi approach for comparative environmental screening, which is crucial to resolving how essential genes contribute to fitness across diverse contexts.

**Figure 3:**
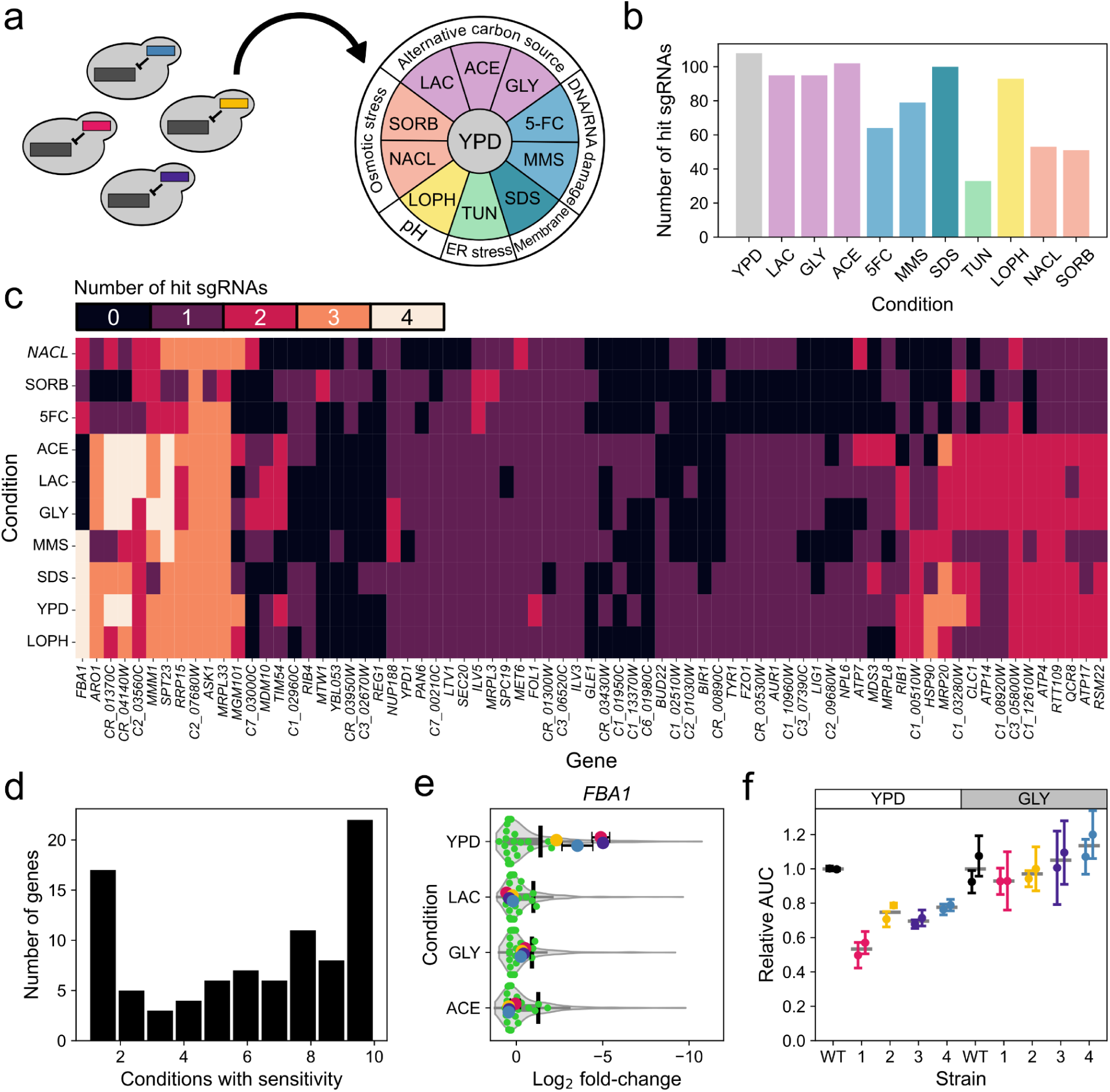
Parallel screening identifies condition-independent repression-sensitive genes. **a)** The same CRISPRi library used in the timecourse assay was screened in 11 conditions in parallel. **b)** Number of significantly depleted sgRNAs in each screening condition. **c)** Number of depleted sgRNAs per gene in each condition, excluding tunicamycin. Only genes which had hits in at least two conditions are shown. **d)** Number of conditions with significant fitness decreases for hit genes in the experiment. **e)** Abundance changes of the 4 sgRNAs targeting *FBA1* in media conditions with (YPD) or without (LAC: YP + lactate 2%, GLY: YP + glycerol 2%, ACE: YP + acetate 2%) a fermentable carbon source. The overall distribution of Log_2_FC for library sgRNAs is shown as a violin plot, and the randomized sgRNAs are shown in green. The black lines show the significance threshold for each condition. **f)** Growth curve assay of strains with *FBA1* targeting sgRNAs in YPD or GLY media. Each measurement was performed in biological duplicates with three technical replicates. Error bars represent the 95% confidence interval around each biological replicate.

From these screens, we identified 23 genes with repression-sensitivity phenotypes that were universal across the 10 environments we analyzed (Figure 3d), representing the most common scenario of effects across conditions. The consistent phenotypes observed for a subset of genes across environments suggest they might be of particular interest for antifungal development. Conversely, some hit genes from the first experiment were found to be repression sensitive only in certain environments. One of the most striking cases was fructose-bisphosphate aldolase *FBA1*, a key enzyme in both glycolysis and gluconeogenesis. In conditions where glucose was the carbon source, *FBA1* showed strong repression sensitivity, which was abolished when switching to non-fermentable nutrients (Figure 3e). We validated this phenotype in small-scale assays, in which growth of *FBA1* CRISPRi mutants was highly impaired relative to the wild type in glucose but not in glycerol media (Figure 3f). Repression efficiency was comparable in both conditions (Figure S8), Because the fungal fructose-bisphosphate aldolase is not homologous to the human isozyme, there are currently efforts to develop specific inhibitors that could be used as antifungals^27^. The ability of *C. albicans* to efficiently perform glycolysis is thought to be important for virulence traits like hypoxia adaptation and biofilm formation^28,29^. On the other hand, the dispensability of *FBA1* on non-fermentable carbon sources suggests that the impact of reduced gluconogenesis flux on fitness is much lower. Thus, the ability to screen CRISPRi strain libraries quickly and efficiently in a large number of experimental conditions can facilitate the detection of environmental effects such as these.

### Many repression sensitivity phenotypes are conserved in antifungal-resistant isolates

In addition to environmental changes, genetic background differences can affect the function and dosage sensitivity of genes^26,30^. This is especially important in the case of antifungal resistance mutations, which can have both positive and negative genetic interactions with other cell components^31–33^. One advantage of pooled library construction and screening is that plasmid libraries can readily be transferred to different genetic backgrounds, enabling parallelized screening and comparative functional genomic analysis (Figure 4a). To validate this approach, we compared the effect of *HSP90* CRISPRi repression in the laboratory strain SC5314 and two clinical isolates, resistant to either fluconazole (FLU^R^)^34^ or caspofungin (CASP^R^)^35^. Both isolates have mutations representative of their respective resistance class: an *ERG11* T299A F449S allele for FLU^R^, and a F641S *FKS1* allele for CASP^R^ ^3^. CRISPRi repression was efficient in both strains (Figure 4b), although its efficiency varied despite no significant changes in baseline *HSP90* transcript levels compared to the wild-type (Figure S9), which might be explained by differences in Cas9 expression^36^. Next, we used the same batch transformation protocol to transform our CRISPRi plasmid library into both drug-resistant strain backgrounds. After sequencing, we found that one transformation batch per background was sufficient to obtain >90% representation of the designed sgRNA, which is higher than what we obtained for the WT strain library (Figure 4c). Overall, this resulted in >80% of individual sgRNAs being present in all three strain libraries and thus directly comparable in downstream assays. Our pooled CRISPRi approach is therefore readily portable, enabling rapid and efficient expansion of large-scale strain libraries across multiple genetic backgrounds, a capability not previously possible in *C. albicans*.

**Figure 4:**
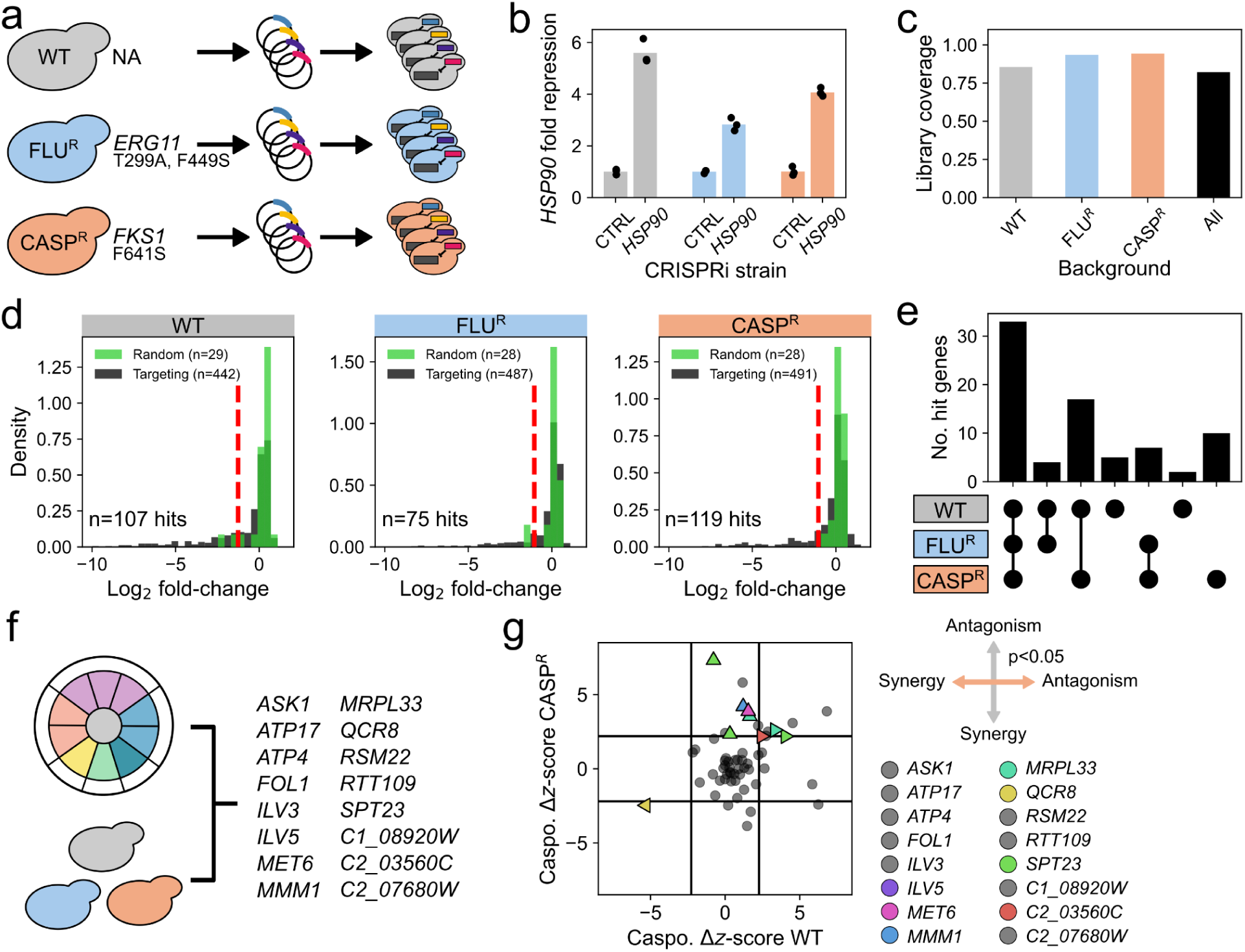
Portable CRISPRi screening of *C. albicans* essential genes identifies a core set of repression-sensitive targets. **a)** The same CRISPRi plasmid library used to build the wild-type strain pooled library was transformed into two new genetic backgrounds, resistant to either fluconazole (FLU^R^) or caspofungin (CASP^R^). **b)** CRISPRi repression of *HSP90* using the same sgRNA in the three different strains. **c)** Library coverage in each genetic background at the start of the screen. The sgRNAs present across all three libraries are shown in black. **d)** Abundance change distribution for each genetic background. The red line indicates the Log_2_FC threshold corresponding to FDR=5%. **e)** Overlap between repression-sensitive genes identified for each strain. **f)** Sensitive genes identified in all environmental conditions and genetic backgrounds. **g)** Interaction between CRISPRi repression and Caspofungin treatment. Only genes with at least one sgRNA showing antagonism or synergy of sufficient magnitude and statistical significance (Benjamini-Hochberg corrected p-value <0.05) are shown as triangles and colored. The direction of the triangle indicates the nature of the interaction, as indicated by the compass (left: synergy in WT, right: antagonism in WT, down: synergy in CASP^R^, up: antagonism in CASP^R^). The black lines show the thresholds for z-score change magnitude (see methods). Genes and sgRNAs not showing synergy or antagonism are shown in grey.

We next used these new libraries to compare the fitness effects of CRISPRi repression in the three different strain backgrounds. We performed competition assays in rich media for the three strains in parallel, and measured relative sgRNA abundance via sequencing. Across all backgrounds, sgRNA abundance changes were highly correlated between replicates, indicating that our pooled screening approach is highly reproducible in clinical isolates (interreplicate correlation for FLU^R^: ρ = 0.69-76, CASP^R^: ρ = 0.85-0.88, p < 0.0001 for all correlations Figure S10). We used the distribution of random sgRNA abundance changes to set a Log_2_FC threshold for each strain (Figure 4d), and identified a similar number of mutant strains with altered abundance in both the wild-type and CASP^R^ strain backgrounds (WT: 107 vs. CASP^R^: 119). We validated our screen’s ability to detect fitness effects in the new strain backgrounds by reconstructing several CRISPRi strains targeting hit genes depleted in both previous experiments and measuring growth curves (Figure S11). As we had observed for the wild-type, the growth curves generally recapitulated the pooled screening results. Among unique genes with at least one CRISPRi fitness defect, we found that 44% (n=34) were represented in all genetic backgrounds (Figure 4e). The second-largest overlap was between the wild-type and the CASP^R^ strain, reflecting the greater sensitivity of the assay in these two backgrounds. We further sequenced the genomes of all three strains to examine whether polymorphisms in promoter regions could have altered sgRNA binding, which could also explain strain-specific fitness effects via reduced or abolished CRISPRi activity. We found that ∼3% of sgRNAs bound promoter regions containing variants unique to clinical isolates (Figure S12), a fraction much too low to explain the differences we observed between strains. These results demonstrate that CRISPRi pooled screening functions reliably across multiple distinct genetic backgrounds, thus extending the ability to perform these high-throughput functional genomic screens to diverse isolates.

Across these pooled CRISPRi screens, we investigated which genes from our set of fungal specific essential genes have repression sensitivity phenotypes, and whether these phenotypes are maintained in different environmental and genetic contexts. Out of the 23 genes that were sensitive to growth in all environments in the wild-type, 16 were also hits in all three genetic backgrounds, allowing us to refine a shortlist of potential new targets to focus on for antifungal development (Figure 4f). As potential new therapeutics might have to be used in combination with existing compounds, understanding whether exposure to antifungals impact repression sensitivity profiles is important. In this scenario, increased CRISPRi fitness defects under treatment could indicate synergy with the drug, whereas reduced fitness defects could indicate antagonism. Ideally, the repression of potential therapeutic targets should be neutral, if not synergistic, to avoid reducing treatment efficiency. The portability and scalability of *C. albicans* CRISPRi pooled screening allow us to examine these relationships while also accounting for genetic background. Thus, we screened the wild-type and CASP^R^ libraries with or without the antifungal caspofungin present at a concentration below the MIC, and compared the effects on sgRNA depletion in both cases. We found that the vast majority of mutant strains showed no changes, with only a few sgRNAs exhibiting large, significant changes in fitness in one but not both strain backgrounds (Figure 4g). For example, one sgRNA targeting *QCR8*, a structural component of respiratory complex III (shown in yellow, lower left quadrant), was more depleted in the wild-type, but did not change significantly in the CASP^R^ background. Only the unsaturated fatty acid synthesis regulator *SPT23* showed a relatively consistent pattern of lower repression sensitivity under caspofungin treatment, with 1 sgRNA in the wild-type and 2 sgRNAs in the resistant strain having reduced effects. We observed a similar pattern with the FLU^R^ strain, with no significant shifts compared to the wild-type (Figure S13). These results suggest that repressing the core repression-sensitive genes we identified should not interfere with at least one major antifungal, which could allow for combinatorial therapy options.

## Discussion

Systematic genetic interrogation of essential genes has transformed our understanding of the core biological processes that regulate microbial life, yet such approaches have remained difficult to implement in fungal pathogens. Here, we establish a regulatable pooled CRISPRi platform for the fungal pathogen *C. albicans* that overcomes the major barriers to high-throughput perturbation of essential genes. By combining regulatable gene repression and rapid construction of high-coverage pooled mutant libraries, our system expands functional genomic capability in *C. albicans* and provides the first application of pooled CRISPRi screening in a fungal pathogen. This new toolkit is available to the community (Addgene #250493) and can be easily used to generate user-specific libraries for diverse functional genomic screening applications in *C. albicans*. Using this platform, we identified a substantial fraction of fungal-specific essential genes that exhibit strong repression sensitivity, assessed core vs. conditionally-essential genes, and demonstrated the portability of this system for comparative functional genomic analysis across diverse *C. albicans* strain backgrounds. Together, our findings underscore the importance of assessing gene essentiality in *C. albicans* as a complex trait that is influenced by gene repression sensitivity, the cellular environment, as well as the genetic context, all of which can impact the evaluation of potential antifungal targets.

Understanding the genetic architecture of essential cellular processes in fungal pathogens remains a longstanding challenge. While foundational studies using deletion libraries and repression libraries^10,16,37^, and transposon mutagenesis^21^ have provided critical early maps of gene essentiality, these approaches rarely capture quantitative or qualitative relationships between gene dosage and fitness. Yet, an accumulating body of work demonstrates that dosage sensitivity is a powerful predictor of druggability^8,9^, and evolutionary constraint in the context of resistance^6^. CRISPRi approaches in the model yeast *S. cerevisiae*^22,38,39^ has highlighted the potential of expression-level perturbations to study critical phenotypes at a high-throughput scale, yet no analogous high-throughput tool existed for major fungal pathogens. By developing a regulatable CRISPRi system compatible with pooled screening, we extend these principles to *C. albicans*, enabling quantitative interrogation of essential gene biology in a critical human fungal pathogen.

An increasing body of research has highlighted the importance of recognizing *C. albicans*, and other fungal pathogen species more broadly, as genetically diverse species in which diverse isolates differ markedly in genome structure, regulatory circuitry, stress tolerance, and drug susceptibility^40–43^. These differences can fundamentally alter gene essentiality and genetic-interaction networks, yet most functional genomics resources in *C. albicans* remain restricted to a single laboratory background. Our findings demonstrate the utility of the CRISPRi system developed here as a highly portable functional genomic tool, enabling scalable, and comparative functional studies across the natural diversity of *C. albicans* strains. By introducing pooled CRISPRi libraries into diverse *C. albicans* strains, including clinical isolates and drug-resistant strains, we provide a route to systematically map background-specific repression sensitivity and identify genetic interactions that are uniquely found in particular genomic contexts. In particular, such analysis can allow us to identify cell components that can alter drug sensitivity uniquely in drug-resistant genetic backgrounds. Previous studies have found that repressing factors such as *PKC1*, *GIN4*, or *CCT8* sensitizes a drug-resistant *fks1\** mutant to echinocandins^44^, illustrating the power of this type of approach at a small scale. Our system now extends this capacity to a large scale, allowing scalable comparative analysis across drug-resistant fungal strains. Integrating genetically diverse isolates into CRISPRi screens will therefore be essential for uncovering strain-specific vulnerabilities, and ultimately designing antifungal strategies robust across clinically relevant genetic backgrounds.

One advantage of our optimized *C. albicans* CRISPRi pooled screening system is its ability to identify gene repression sensitivity at scale, thereby enabling the systematic identification of essential genes for which even modest reductions in expression lead to substantial growth defects. These highly repression-sensitive genes represent particularly compelling antifungal targets, as therapeutic inhibition may require only partial suppression of gene function, thus broadening the range of compounds that can be effective therapeutics at low doses, and reducing the risk of off-target toxicity. By identifying fungal-specific, repression-sensitive genes that are essential for fungal viability across diverse environmental and genetic contexts, our study generates a prioritized list of targets for antifungal drug discovery. This resource coincides with emerging AI-driven drug discovery pipelines, including protein structure-prediction tools, generative molecular design, and machine-learning programs that identify ligandable pockets and predict inhibitor binding and affinity^45–47^. As protein structures for many of the identified targets can now be rapidly modelled, AI-enhanced screening can accelerate the development of small molecules tailored explicitly to these fungal vulnerabilities. Thus, our CRISPRi-based platform provides a bridge between computational drug design efforts and the antifungal drug development pipeline.

Together, we present a fast, scalable, and portable CRISPRi system that enables high-throughput functional genomics in *C. albicans*, including multi-environment screening, comparative analysis across diverse genetic backgrounds, and interrogation of repression-sensitive essential genes. This platform provides the first pooled CRISPRi screening framework for a major fungal pathogen and reveals a core set of fungal-specific, repression-sensitive essential genes with broad relevance for antifungal target discovery. Given its flexibility and efficiency, this system may now be adapted to genome-scale libraries, additional fungal species, and a wide range of biological questions, thus providing an important new strategy for dissecting fungal biology.

## Materials and methods

Strains, plasmids and oligonucleotides used in this study are shown in Table S2, S3 and S4, respectively. Media recipes are available in Table S5.

### Regulatable CRISPRi expression plasmid construction

To create the regulatable CRISPRi plasmid, the tetO promoter sequence and tetracycline transactivator (TAR) were synthesized de novo by Synbio Technologies LLC and inserted into the original CRISPRi plasmid in place of the original ACT1 promoter driving expression of *dCas9*. Briefly, the backbone of the plasmid was taken from the original CRISPRi plasmid (pRS159, Addgene # 122378) previously created by the Shapiro lab^14^. The tetO promoter and TAR were synthesized using a *C. albicans* codon-optimized plasmid provided by the Cowen lab (pRS250, Cowen lab reference pLC1031) as the sequence reference. The *ACT1* promoter that previously expressed dCas9 in pRS159 was removed and the TAR and promoter amplicon was cloned in its place. The plasmid was transformed into *E. coli* DH5α cells and grown at 30 °C in LB media containing both ampicillin (AMP) and nourseothricin (NAT). This plasmid has been deposited on Addgene (on hold, #250493).

### sgRNA library design

The subset of essential genes we selected for this library was based on data from a recent TN-Seq assay in a haploid *C. albicans* strain^21^. Briefly, after filtering based on putative essentiality (as measured by haploid TN-Seq data), absence of human homologs and conservation in three other distant fungal pathogens (*Aspergillus fumigatus*, *Histoplasma gondii*, and *Cryptococcus neoformans*), the authors identified a set of 130 fungal specific conserved essential genes. SgRNA design was performed broadly as described, with some minor modifications to adjust for the pooled library format^15^. For each of these genes, we searched for potential target sites in their promoter region (up to 300 bp upstream from the start codon) with the EuPaGDT online tool and used the predicted score to select the best four sgRNAs. Promoter sequences were retrieved from the *Candida* Genome Database (CGD^48^). Target sequences that contained Sap1 or Pac1 recognition sites were filtered out, as these enzymes are used in subsequent cloning steps. As a negative control, we also generated four sgRNAs targeting the well-characterized molecular chaperone, *Hsp90*, as a positive control and 30 randomly randomized guide sequences with no perfect matches in the *C. albicans* genome as a negative control. This resulted in a total of 554 guides in the library. The oligonucleotide sequences were designed as follows:

5’ GGTCTAATTT**GCTCTTCC**cga-N20-gtt**GGAAGAGC**TTTTAATTAAGCTT 3’ (bold: SapI recognition sequences, lowercase: ligation overhang for directional cloning, N20: 20 bp target sequence). The resulting oligonucleotide library was then ordered from IDT as pooled single-stranded DNA (ssDNA).

### Primer extension

Pooled ssDNA sequences were converted to double-stranded DNA by PCR. Two PCR reactions were set up each with a 50 μL total volume. Each reaction consisted of 1 μL (1 μg/μL) of ssDNA (oligonucleotide pool), 1 μL of primer (pRS 596) 5’ AAGCTTAATTAAAAGCTCTTC 3’ (equimolar amount of primer, 318.18 ng/μL), 41 μL of Nuclease-free water and 5 μL of 10x NEBuffer 2.1 (NEB, cat. B7202S). The PCR conditions were set to: 94 °C for 5 mins, slow ramp down (0.1 °C/sec) to 46.4 °C and hold for 5 mins, slow ramp down to 37 °C and hold for 1 min. Then, 1 μL of 5,000 U/mL DNA polymerase I, Large (Klenow) fragment (NEB, cat. M0210S) and 1 μl of 10 mM dNTPs (NEB, cat. N0447L) was added to the tube. The thermocycler conditions were then set to 37 °C for 1 hr, followed by a heat-inactivation step at 72 °C for 20 mins, a slow ramp down to 42 °C for 1 min, and then held at 4 °C until ready to use. The 50 μL reaction was then processed using the DNA Clean and Concentrator-5 kit (Zymo research, cat. D4013) as per manufacturer’s instructions. The cleaned product was quantified using a Tecan Infinite Pro with a NanoQuant plate.

### Golden Gate cloning

The double-stranded oligonucleotide pool (dsN20) sequences were cloned into the inducible CRISPRi plasmid using a Golden Gate cloning strategy previously described^14,15^. Briefly, the following reaction was prepared in quadruplicate: 10 μl of miniprepped CRISPRi plasmid (180 ng/μl), 1 μl of dsN20 (diluted to a concentration that results in a 3:1 insert to vector ratio), 2 μl 10mM ATP (NEB, cat. P0756S), 2 μl 10x rCutsmart buffer (NEB, cat. B6004S), 1 μl 400,000 U/mL T4 DNA ligase (NEB, M0202S), 1 μl of 10,000 U/mL Sap1 (NEB, cat. R0569L) and 3 μl nuclease-free water. The thermocycler was then set for the following conditions: 37 °C for 2 mins, 16 °C for 5 mins, cycled 99x, 65 °C for 15 mins, 80 °C for 15 mins.

### Bacterial transformation

Before transforming, the cloned plasmids were first treated with 1 μL of SapI and incubated for 1 hr at 37 °C to linearize any remaining uncloned plasmids. Next, we transformed the cloned library into *E. coli* DH5α competent cells using a standard chemical transformation: 5 μl of cloned plasmid was added to 50 μL of commercial chemically competent cells (NEB, cat. C2987H) and incubated on ice for 30 mins, heat shocked at 42 °C for 30 secs, and then incubated on ice for an additional 5 mins. After this, 950 μL of SOC media (NEB, cat. B9020S) was added. The cells were then incubated horizontally shaking for 1 hr and 30 mins at 30 °C. From the tubes, 50 μl of culture and 150 μl of water were plated on LB AMP (100 μg/mL, BioShop, cat. AMP201.25) and NAT (50 μg/mL, Jena Bioscience, cat. AB-102L) plates to collect information on colony counts. To prepare the LB-semi-solid media, 0.35% seaprep agar (Lonza, cat. 50302) and LB media mix (BioShop, cat. LBL407.500) was autoclaved with the stir bar left in. The media was cooled to room temperature at which point AMP at a concentration of 100 μg/mL and NAT at a concentration of 50 μg/mL was added to it. As per the methods from Momen-Roknabadi *et al* 2020^22^, 1 L of semi-solid agar can hold between ∼1-5 million colonies and therefore media should be prepared accordingly for the amount of transformation and the efficiency of the transformations^22^. The remaining transformation culture was added to the semi-solid agar and stirred for 10 mins. The flask was then put in an ice water bath where the water level was above the media line in the flask for 1 hr to solidify. The flask was then carefully moved to a 30 °C static incubator and left to grow for 24-36 hrs. From the plates, 30 individual colonies were sequenced using Sanger sequencing as a quality control check: all clones had a properly cloned sgRNA and did not contain any empty backbone plasmids. Cells were then harvested from the agar by heating the media at 37 °C for 20 mins while stirring with the magnetic stir bar to break apart the agar and then centrifuged to collect a pellet which was then resuspended in 5-10 mL of LB-AMP-NAT. The pool was then frozen in 2 mL aliquots by combining 1 mL of culture (at an OD_600_ of ∼30) with 1 mL of 40% glycerol.

### Plasmid pool extraction

A maxi prep plasmid extraction (Nucleobond Xtra Maxi EF, Macherey-Nagel, cat. 740424.10) was performed to obtain a large quantity of plasmid DNA to be used in the fungal transformation protocol. To prepare the culture for extraction, 600 mL of LB, AMP (100 μg/mL) and NAT (50 μg/mL) was added to a 2 L flask. We then added 1 aliquot of the frozen bacteria pool to the media, and cells were grown at 30 °C for 12-18 hours until an OD of > 2 was reached. Extraction was then performed as per manufacturer’s instructions. The extracted plasmids were quantified using a Tecan Infinite Pro with a NanoQuant plate.

### *Candida albicans* transformation

Plasmids were transformed into *C. albicans* using the LiAc-PEG chemical transformation protocol, as previously described^15,49^. A single transformation batch comprises 14 individual transformations, which are then subsequently pooled. Briefly, 30 μL of plasmid DNA (∼225 ng/μL) was combined with 1 μl of 10,000 U/mL PacI (NEB, cat. R0547L), 5 μL of 10x rCutsmart and 14 μL of Nuclease-free water and was digested overnight at 37 °C static. A 50 mL overnight culture of the *C. albicans* strain of interest was grown at 30 °C with shaking (250 rpm) for roughly 16-18 hours. From the overnight culture, calculate the volume of culture needed to obtain roughly ∼5×10^7^ cells. Pellet cells using centrifugation and remove the supernatant. Prepare a transformation master mix using the following recipe: 800 μl of 50% polyethylene glycol (PEG), 100 μl of 1 M lithium acetate (pH 7.4) 100 μl of 10× Tris-EDTA (TE) buffer, 40 μl of 10 mg/mL ssDNA (salmon sperm, ThermoFisher cat. 15632-011), and 20 μL of 1 M dithiothreitol (DTT). The 1060 μl transformation mix and 45 μl (∼6000 ng of linearized plasmid) were then added to a cell pellet, and the cells were flicked to resuspend them. The cells were incubated static at 30 °C for 1 hr and then heat-shocked at 42 °C for 45 mins, before being washed twice with YPD and resuspended in 5 mL of YPD with ATc (250 ng/mL) media for 4 hr at 30 °C with shaking (250 rpm). After outgrowth, these cells were then pelleted and resuspended in 300 µL of YPD added to 1 L of YPD-semi-solid agar (0.35% seaprep agar) containing 250 ng/mL of ATc (Alfa Aesar, cat. J66688) and 250 µg/ml NAT. To get a rough estimation of the transformation efficiency 50 µL from the transformation culture was plated on YPD-NAT-ATc plates and grown at 30 °C static for 48 hours. The 1 L of semi-solid agar was divided between three fernbach flasks as we noticed that *C. albicans* cells grew poorly past a ∼1.5 cm depth in the agar. Flasks were grown at 30 °C for 48-72 hours (to a point at which colonies became visible). The cells were then recovered using the same method as described for the bacterial transformation, pelleted, and frozen at −80 °C in 2 mL aliquots containing 1 mL of pooled library and 1 mL of 40% glycerol.

### Time-course competitive growth assay

A timecourse assay was used to monitor the change in mutant abundance over time. For this test an aliquot of the wildtype fungal pool was thawed on ice for 15 mins and then added to a 250 mL flask containing 50 mL of YPD and ATc (final concentration 250 ng/mL). The flask was incubated shaking (200 rpm) at 30 °C for 12 hrs. The flask was then removed from the incubator and a subculture was prepared to an OD_600_ of 0.05 in 50 mL of YPD and incubated shaking (200 rpm) at 30 °C for 12 hrs. The culture was then removed from the incubator and a 1 mL sample of the culture was labeled timepoint 0 (TP0) and frozen at −80 °C. To three 50 mL falcon tubes 15 mL of YPD was added and to another three tubes 15 mL YPD+ATc (250 ng/mL) was added. The TP0 culture was subcultured into these tubes to an OD_600_ of 0.05 and incubated shaking (200 rpm) at 30 °C for 12 hrs. The six tubes were then subcultured again in their respective media types and grown for 12 hrs under the same conditions. This was then repeated for a third and final passage. At the end of each 12 hr out growth period (passages), 1 mL samples of each culture were frozen at −80 °C.

### Multi-environmental condition competitive growth assay

For this test an aliquot of the wildtype fungal pool was thawed on ice for 15 mins. Three 50 mL falcon tubes were labeled TP0-YPD (time-Point 0) replicates (R)1-3 and were filled with 10 mL of YPD. Another three falcon tubes were labeled TP0-ATC R1-3 and were filled with 10 mL of YPD+ATc (250 ng/mL). To each tube, 200 µL of the thawed pool was added. The cultures were incubated for 12 hrs at 37 °C shaking (250rpm). These cultures will be used as timepoint zero to determine the initial frequency of each mutant in the pool. Media was prepared for each of the conditions of interest (see Table S5 for media recipes). Media conditions were prepared with and without ATc. To a 24 deep well plate (Fisher Scientific, cat. 14-222-349), 3 mL of each condition (with and without ATc) were added to three wells. The TP0 cultures were subcultured into each well at an OD_600_ of 0.1 with TP0-YPD R1 being subcultured into the well labeled R1 for each condition, following this for each of the other replicates and conditions. The subcultures were grown at 37 °C shaking (550 rpm) for 12 hrs. After the first growth period the subculturing method was repeated for another two rounds of passaging. After the third passage’s outgrowth was completed, samples were taken from each of the replicates. Sample volumes contained at least ∼ 5×10^7^ cells and were frozen at −80 °C.

### Competitive growth assay to evaluate changes in abundance in different genetic backgrounds

For this test an aliquot each of the three fungal pools was thawed on ice for 15 mins. For each fungal pool three 50 mL falcon tubes were labeled with the respective genetic background name and TP0-YPD (time-Point 0) replicates (R)1-3 and were filled with 10 mL of YPD. Another three falcon tubes were labeled TP0-ATC R1-3 and were filled with 10 mL of YPD+ATc (250 ng/mL). To each tube, 200 µL of the corresponding thawed pool was added. The cultures were incubated for 12 hours at 37 °C shaking (250rpm). These cultures will be used as timepoint zero to determine the initial frequency of each mutant in the pool. To a 24 deep well plate, 3 mL of YPD were added to three wells for each fungal pool and 3 mL of YPD+ATc were added to another three wells for each fungal pool. The TP0 cultures were subcultured into each well at an OD_600_ of 0.1 with TP0-YPD R1 being subcultured into the well labeled YPD R1, following this for each of the other replicates and pools. The subcultures were grown at 37 °C shaking (550 rpm) for 12 hrs. After the 12 hrs the subculturing method was repeated for another two rounds of passaging. After the third passage’s growth period was completed, samples were taken from each of the replicates. Sample volumes contained at least ∼5×10^7^ cells and were frozen at −80 °C.

### Genomic DNA extraction

Genomic extractions were performed using the PureLink™ Genomic DNA Mini Kit (Invitrogen, cat. K182001) or with the phenol-chloroform method^50^. With the kit, frozen samples were thawed, spun down, and the supernatant was removed. A master mix was prepared as follows: 4.55 g of D-Sorbitol (BioShop, cat. SOR508.1), 500 μL of 0.5 M EDTA (VWR, cat. E177-500ML), 0.5 M, 25 μL of β-mercaptoethanol (BioShop, cat. MER002.100), 0.0335 g Zymolyase (22,400 units/g, BioShop, cat. ZYM001.1), 24.48 mL nuclease-free water (final concentration of master mix 1 M sorbitol, 10 mM EDTA, 14 mM β-mercaptoethanol, 15 units/500 μL zymolyase). The cell pellet was resuspended in 500 μL of the master mix and incubated at 37 °C for 1-2 hrs (for cells from the clinical isolates backgrounds a longer incubation of 2 hrs was used). After incubation manufacturing instructions were followed to perform the kit genomic purification. After samples were extracted and purified, DNA concentrations were determined by Qubit analysis (analysed using a Qubit Flex Fluorometer with a high sensitivity DNA kit cat. Q33230) or by absorbance using a Tecan Infinite Pro with a NanoQuant plate, following the manufacturer’s instructions.

For samples with low yields a phenol chloroform method was used in place of the kit. Briefly, frozen pellets were thawed and spun down into pellets and resuspended in 200 μL a homemade lysis buffer (2% Triton X-100, BioShop, cat. TRX777.500), 1% SDS, 100 mM NaCl, 10 mM Tris-Cl (pH 8.0), 1 mM Na2EDTA). To the resuspended sample 200 μL of phenol:chloroform:isoamyl alcohol (25:24:1) and a scoop (roughly the size of the original cell pellet) of acid washed glass beads (Sigma Aldrich, cat. G8772-100G) were added to each sample. Samples were beaten in a bullet blender for 4 mins. Samples were then pelleted at 13,000 xg for 5 mins. The top (aqueous) layer was transferred into a new “tough” micro-centrifuge tube (Sigma, cat. EP022363344). To the separate aqueous layer, 200 μL of phenol:chloroform:isoamyl alcohol (25:24:1) was added and the mixture was vortexed for 1 min. Samples were then pelleted at 13,000 xg for 5 mins at room temperature. The top (aqueous) layer was transferred to a standard microcentrifuge tube. A 1/10 volume of 3 M pH 5.2 sodium acetate, and 2 volumes of chilled 95% ethanol was added to the tube and pipetted to mix. Samples were then incubated at −20 °C for 15 mins. Samples were then centrifuged at 13,000 xg for 15 mins at 4 °C. The supernatant was then discarded. The pellet was then re-suspended in 400 μL of pH 8.0 TE buffer with 3 μL of 10 mg/mL RNase and incubated for 5 mins at 37 °C. After which 40 μL of 3 M pH 5.2 sodium acetate, and 800 μL of pre-chilled 95% ethanol and incubated at −20 °C for 15 mins. Samples were then centrifuged at 13,000 xg for 15 mins and the supernatant was discarded. The pellets were then dried in a 37 °C incubator for ∼10 mins. Pelleted DNA was then re-suspended in 50 μL of Tris-HCl (pH 8.0) or TE buffer (pH 8.0). Nanodrop was then used to quantify the gDNA.

### NGS library preparation and sequencing

NGS library preparation was performed in two successive PCR reactions. The first PCR involved the use of amplicon specific primers attached to illumina sequencing adaptors. These primers were designed to amplify the sgRNA region in the plasmid. Genomic DNA was diluted to 30 ng/μL. Briefly a master mix was prepared with 25μl of 2x NEBnext (NEB, cat. M0541L) and 5 μL of primers (see Table S4) diluted to 10 μM concentration per reaction. To a PCR tube 30 μL of master mix and 20 μL diluted genomic DNA was added to PCR tubes for each reaction and then run under the following thermocycler conditions 98°C for 30 sec, cycle the following three parameters 25 times: 98 °C for 10 sec, 61 °C for 30 sec, 72 °C for 30 sec, then a final step at 72°C for 2 mins. Following the PCR, each reaction was cleaned using the NucleoMag clean up kit (Macherey-Nagel via D-MARK Biosciences, cat 744970.5) as per manufacturers instructions using a 1.6:1 bead to DNA ratio. The clean-up resulted in 30 μL of cleaned amplicon product. Qubit analysis was then used to quantify amplicon concentrations. For the second PCR, a set of illumina index primers (see Table S4) were used. For this PCR a single reaction was set up for each sample. A mastermix is prepared by adding 25 μL of 2x NEBnext and 15 μL of nuclease-free water for each reaction. To each PCR tube, 40 μL of the master mix is added with 5 μL of 5 μM illumina index primer pairs and 5 μL of the cleaned amplicon (diluted to 3 ng/μL). The following thermocycler conditions were then used: 98 °C for 30 sec, cycle the following three parameters for 10 cycles: 98 °C for 10 sec, 55 °C for 30 sec, 72 °C for 30 sec, then a final step at 72 °C for 2 mins. From the reaction 45 μl of the PCR reactions was then cleaned using the Nucleomag clean up kit with a bead to DNA ratio of 1:1. The final elution step resulted in 35 μL of cleaned amplicon product. Qubit analysis was then performed on the samples to obtain the DNA concentration of each sample. Samples were diluted to 5 ng/μL and were sent to the University of Guelph Advanced Analysis center for quality control through RT-qPCR analysis to identify if indices were correctly incorporated into the amplicon sequences. Libraries were sequenced on either Illumina MiSeq or NovaSeq platforms. Samples were run on an Illumina NovaSeq 6000 with a S4 flow cell for 150 cycles paired-end reads.

### Competition assay data analysis

All data analysis and visualization was performed in Python 3 unless stated otherwise, using the following packages: numpy v1.26.4^51^, matplotlib v3.9.2^52^, scipy 1.13.1^53^, pandas v2.2.2^54^, seaborn v0.13.2^55^. Forward and reverse reads were first assembled using Pear v0.9.11^56^, adjusting overlap parameters depending on read length. The merged reads were then trimmed to keep only a 80 bp region centered on the sgRNA, reverse complemented, and dereplicated using Fastx-toolkit v0.0.14^57^, keeping the counts associated with each unique sequence. These unique sequence clusters were then associated with library sgRNAs by perfect match of the sgRNA sequence and an 8-nucleotide anchor on each side to obtain read counts. Counts were normalized into frequencies using the following formula:

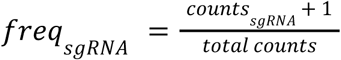

From there, we measured changes in relative abundance using log_2_-fold change:

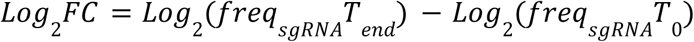

We used a minimal read count threshold of 50 reads at T0 to filter out sgRNAs that were poorly covered. We used the median of Log_2_FC values across replicates for downstream analysis. To determine what constituted a significant change in abundance, we fit a normal distribution to the Log_2_FC values of the randomized sgRNAs. This allowed us to predict the expected false positive rate as a function of Log_2_FC values, and from there we set the significance threshold as the Log_2_FC value at which the false discovery rate reached 5%. One of the random sgRNAs (#18, AGCTCTACTGTTACAAGGGG) was found to induce CRISPRi specific fitness defects reproducible across multiple screening conditions and genetic backgrounds, and as such was excluded from the control distribution. GO annotations were retrieved from the *Candida* Genome Database^58^. Thresholds were set separately by environmental condition, genetic background, or background by environment combination. For Figure 4g and Figure S12, we normalized the Log_2_FC values of each condition using the parameters of the randomized sgRNA distribution of each condition. The threshold for a significant difference in z-score under caspofungin exposure was set to the mean of the normalized abundance change thresholds at 5% FDRwith or without caspofungin 5% FDR.

### Growth curve assays

Overnight cultures of *C. albicans* grown in YPD were diluted to an OD_600_ of 0.01 in YPD. From this dilution 100 μL was added to a 96 well plate containing 100 μL of YPD to obtain a final volume of 200 μL. Growth conditions were set to 37 °C with orbital shaking for 1000 sec at a 4 mm amplitude. Optical density measurements were taken at 20 min intervals over the course of 24 hours using a Tecan Infinite 200 PRO microplate reader.

### RT-qPCR

Differential gene expression within mutants was performed through RNA-extractions and RT-qPCR. Overnight cultures of *C. albicans* were subcultured to an OD_600_ of 0.05 in 40 mL of YPD and grown at 37 °C shaking at 250 rpm to an OD_600_ between 0.2-0.5. Cultures were then centrifuged at 4000 rpm for 10 mins and the pellet was resuspended in ∼ 200 μL of YPD and frozen at −80 °C. RNA extractions were performed using PuroSPIN™ Total RNA Purification Plus Kit (Luna Nanotech, cat. NK251-200). RT-qPCR was performed using a One-Step RT-qPCR kit (NEB, cat. E3005L). Real-time PCR was performed using a QuantStudio 3 Real-Time PCR system from Thermo Fisher Scientific Inc under the following cycling conditions 55 °C for 10 mins, 95 °C for 1 min, 40 cycles of a two-step qPCR (95°C for 10 sec and 60 °C for 30 sec) and a Melt curve conditions were set for 95°C for 15 s, 60°C for 1 min and 95°C for 1 sec. The C_T_ values for the experimental gene of interest were compared to the housekeeping gene to obtain a Δ*C_T_* value for each strain. The Δ*C_T_* for the mutants were compared to the Δ*C_T_* of the control strain to obtain a ΔΔ*C_T_* value and used to calculate a fold difference in gene expression of the experimental gene of interest.

### Whole Genome Sequencing (WGS)

Genomic extractions were performed using the Genomic extraction protocol described above, and samples were sent to Genscript for sequencing. WGS sequencing data was analyzed using a modified version of the Broad Institute Funpipe workflow with GATK v4.6.1.0^59^, Picard v3.1.0^60^, bwa199 v0.7.18^61^, samtools v1.20^61^, bcftools v1.19^62^, bedtools v2.31.0^63^. Briefly, adapter sequences were trimmed using GATK and Picard, and reads were aligned to the C. albicans SC5314 haploid reference genome (ASM18296v3203) using bwa-mem with default parameters. The alignments were then processed with GATK to prepare for variant calling using Haplotype caller with default settings except for -ploidy = 2. Variant calls were filtered using the following thresholds: for SNPs, QD > 12.5, QUAL > 30, SOR < 3, FS < 60, MQ > 40, and for indels QD > 20, QUAL > 30, FS < 200. After hard filtering, the variants were cross-referenced with a list of known heterozygous positions from a phased assembly^58^ and with sgRNA target sequence genome mapping generated using bowtie^64^. Potential variant sites were examined manually using IGV^65^. We obtained high coverage for all three strains (WT: 83x coverage, ARB: 72x coverage, ERB: 71x coverage). No cases of aneuploidy, which are sometimes observed in antifungal-resistant strains, were detected. For the azole-resistant strain ARB, the presence of homozygous missense mutations T299A and F449S in *ERG11*, which are known to be sufficient to confer fluconazole resistance were validated. In ERB, the presence of a homozygous echinocandin resistance mutation (F641S) in *GSC1* (also known as *FKS1*) was validated.

## Data availability

The sequencing data for pooled competition assays and whole-genome sequencing was deposited on the NCBI SRA (accession number:PRJNA1381690). Read counts and Log_2_FC values of sgRNAs for each screen are provided as supplementary Tables S6, S7 and S8.

## Code availability

Data analysis was performed using custom python 3.12.7 notebooks and scripts, available at https://github.com/TheShapiroLab/Calbicans_Pooled_CRISPRi_2025/.

## Acknowledgements

This research was enabled in part by support provided by SHARCNET (sharcnet.ca), Calcul Québec (calculquebec.ca) and the Digital Research Alliance of Canada (alliancecan.ca). The authors thank C. Landry for comments on the manuscript.

## Supplementary Figures 1-13

**Figure S1:**
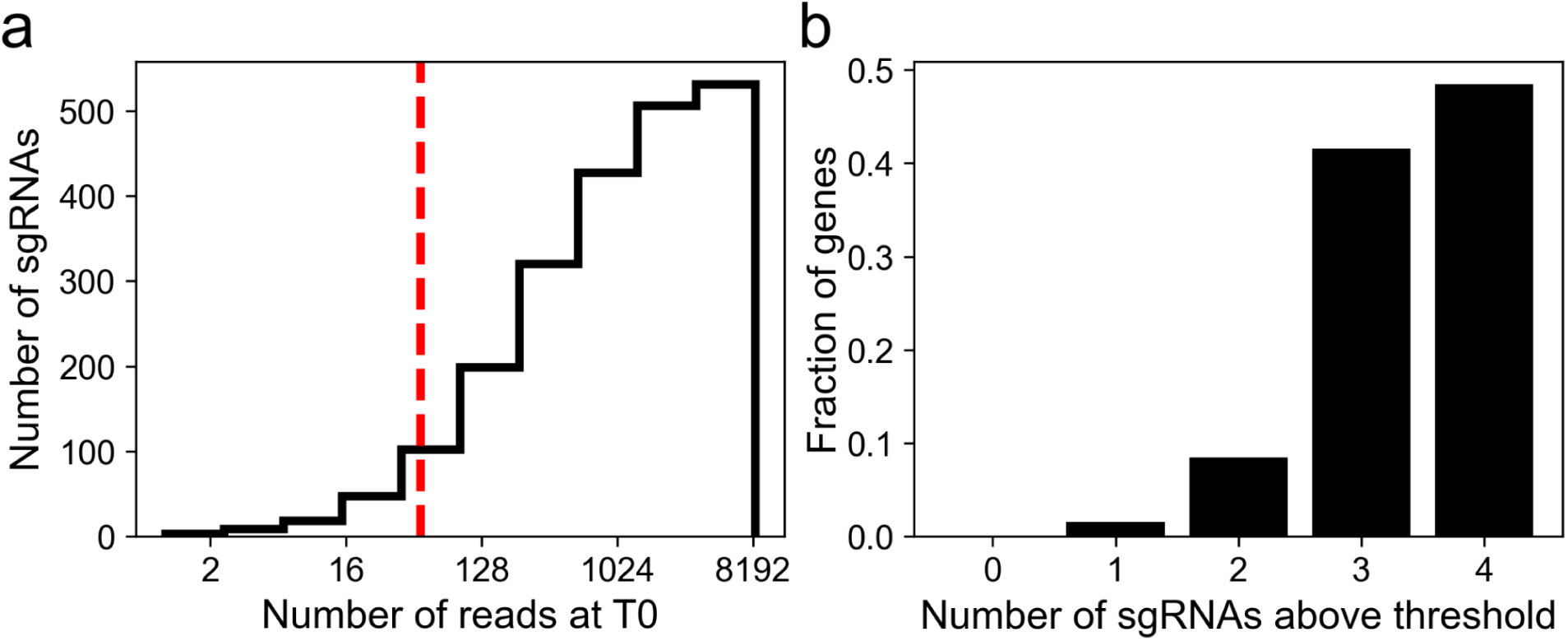
Library coverage in the *C. albicans* wildtype strain pool. **a)** Initial coverage distribution in the library for the 551 sgRNAs. The red dotted line represents the minimal T0 abundance threshold for inclusion in Log_2_FC calculations, 50 read counts. **b)** Proportion of genes with partial or full sgRNA representation in the library.

**Figure S2:**
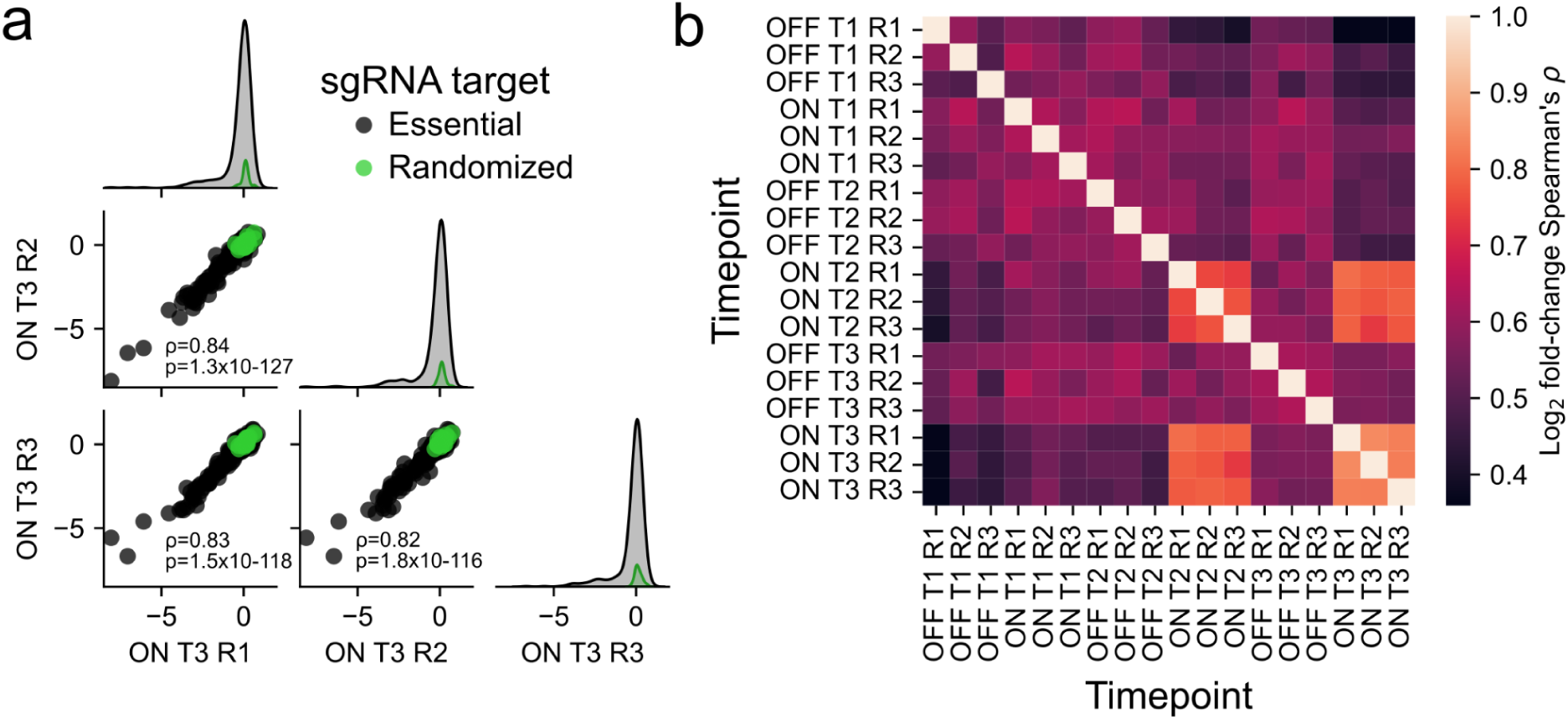
Replicate correlation in the time-course competition assay. **a)** Log_2_FC in CRISPRi ON conditions after the third passage across replicates. Randomized sgRNAs (n=29) are shown in green, and those targeting essential gene promoters (n=438) are in black. The correlation between values was measured using Spearman’s rank correlation. **b)** Spearman’s rank correlation matrix across all time-course samples (n=469 sgRNAs per comparison).

**Figure S3:**
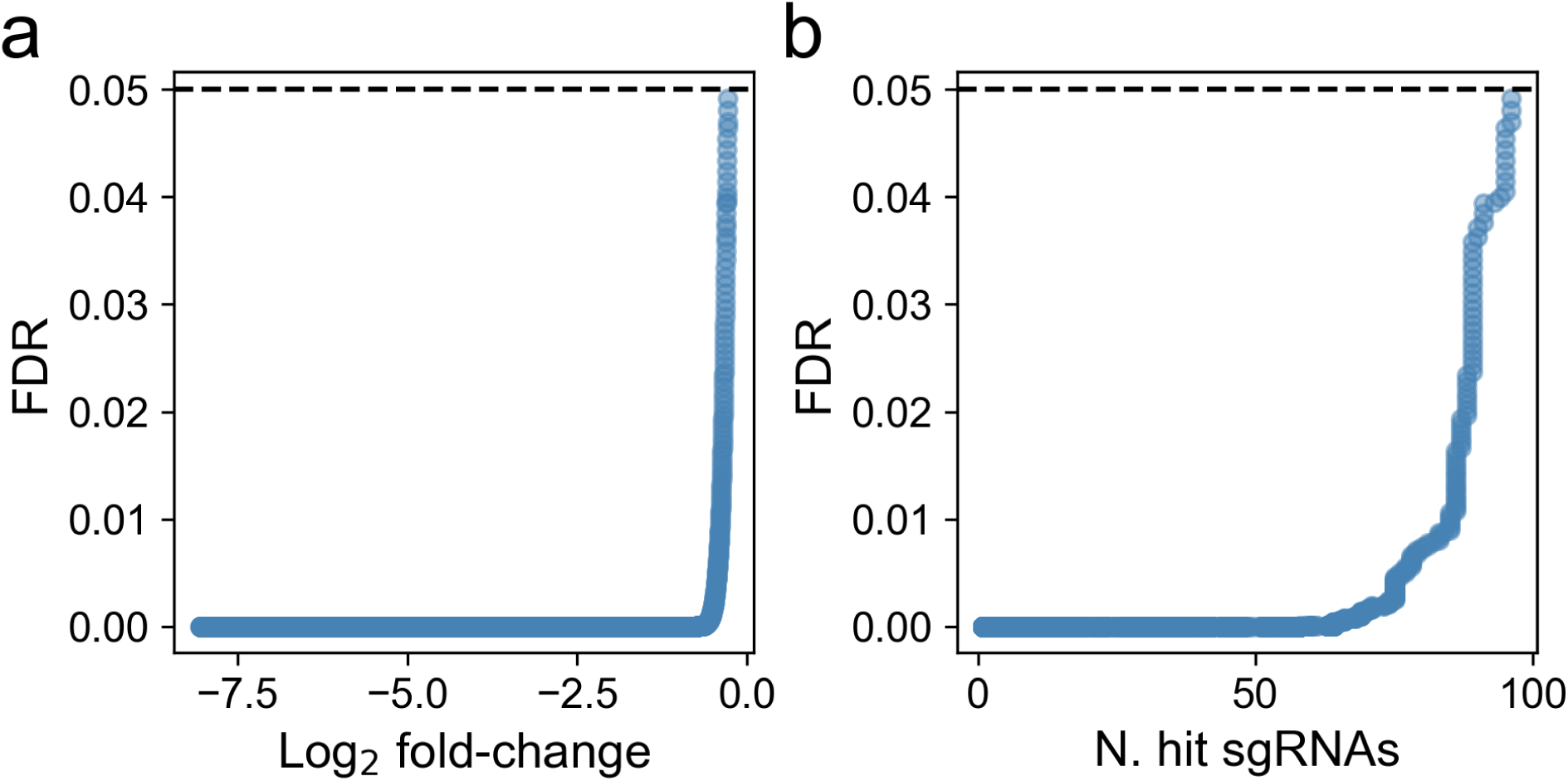
Significance threshold for sgRNA depletion. **a)** False Discovery Rate (FDR) as a function of Log_2_FC threshold, based on the distribution of randomized sgRNAs abundance change and the number of depleted sgRNAs at that threshold. **b)** FDR as a function of the number of sgRNAs determined to be significantly depleted.

**Figure S4:**
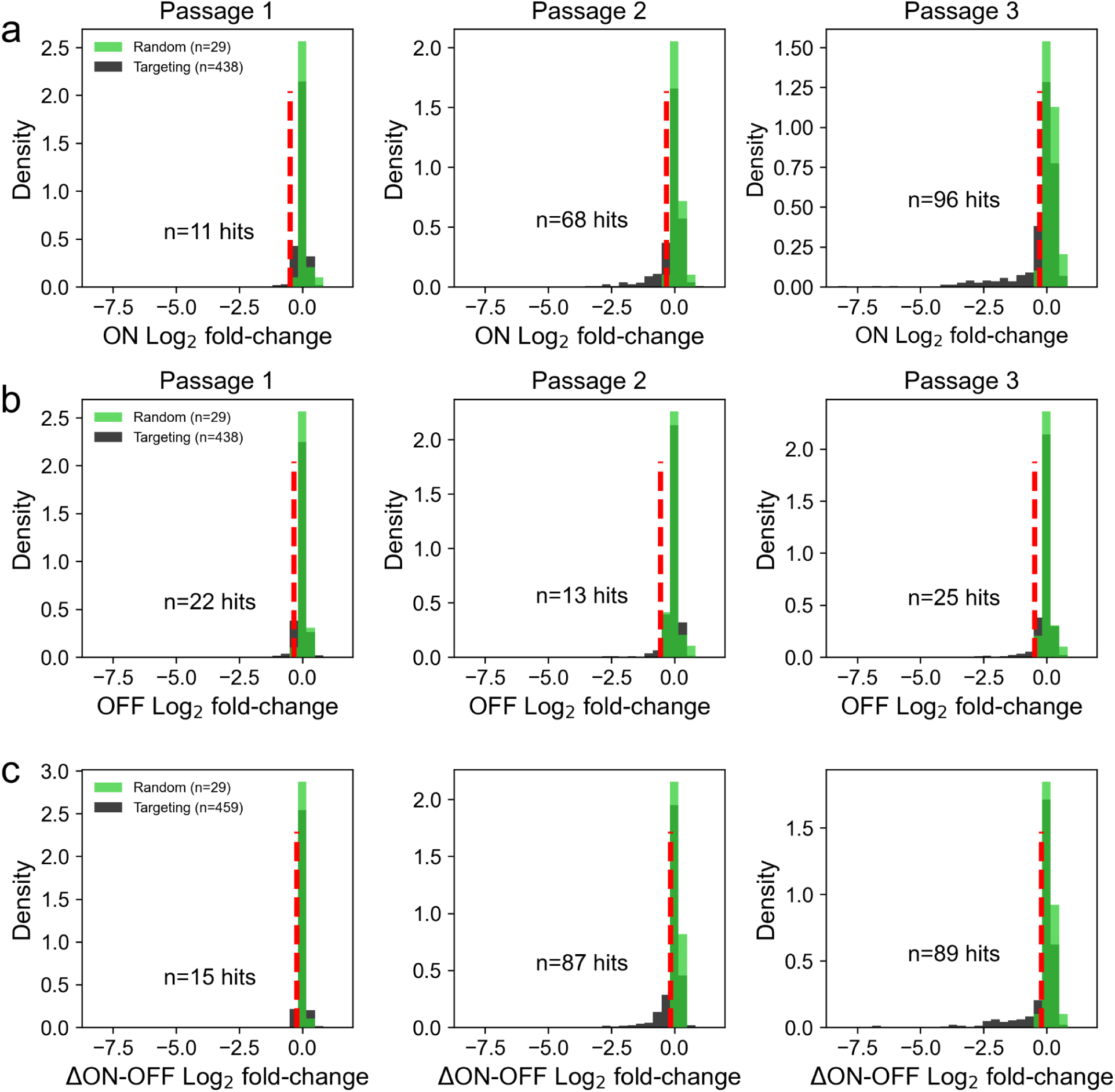
Distribution of abundance changes across passages. In all plots, the dotted red line shows the significance threshold for depletion when comparing library and randomized sgRNAs. **a)** Abundance changes for CRISPRi-ON (media without ATc) samples. **b)** Abundance change for CRISPRi-OFF samples (media with ATc) **c)** Distributions of CRISPRi-ON/OFF Log_2_FC differences for the different passages.

**Figure S5:**
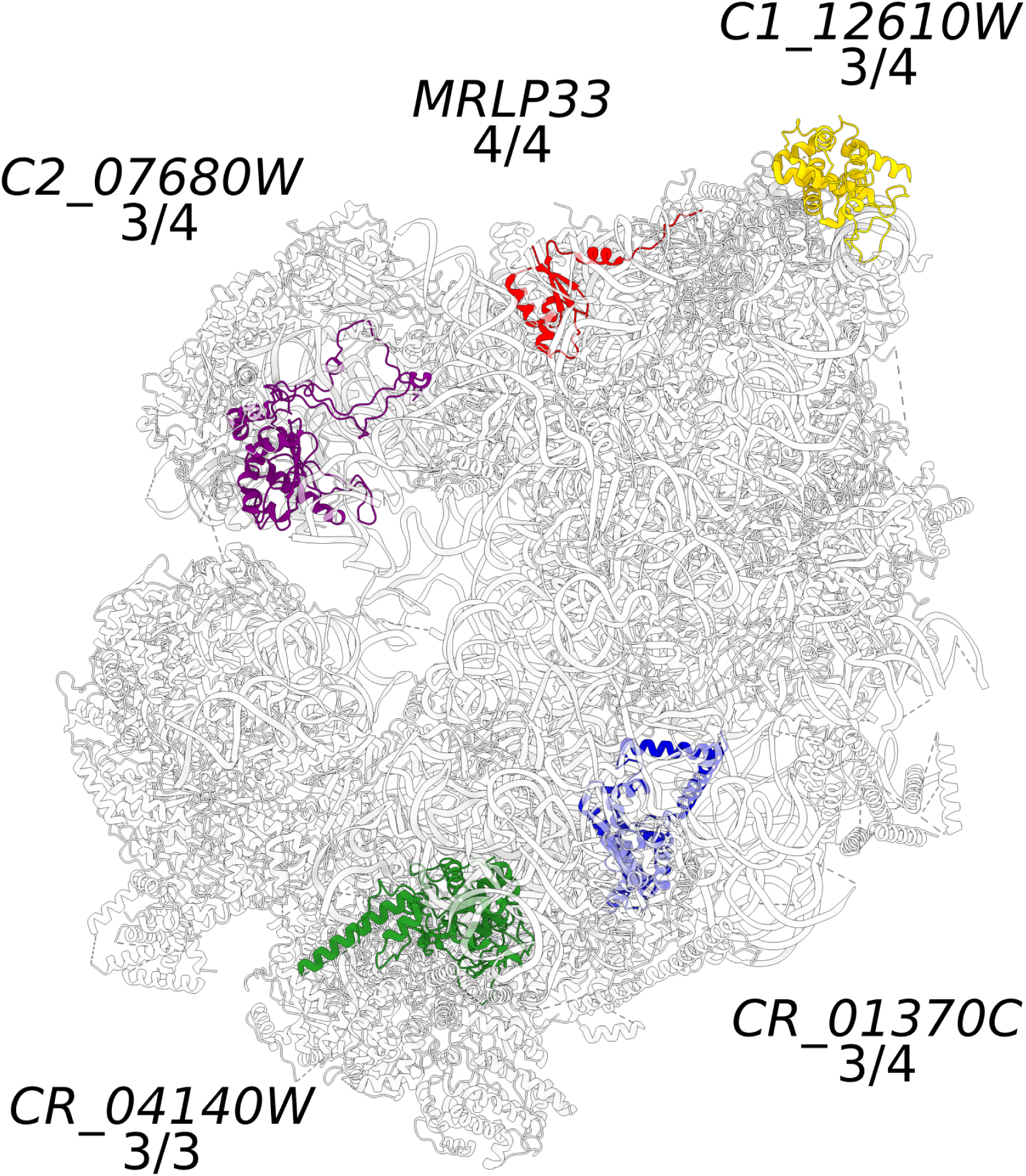
Repression sensitive hit genes encoding mitochondrial ribosomal subunits do not cluster in space. The structure shown is the *Saccharomyces cerevisiae* mitochondrial subunit (pdb: 5MRC ^66^, with orthologs corresponding to putative repression sensitive genes in *C. albicans* : *MRPL33* (*S.c. MRPL33*, red)*, C1_12610W* (*S.c.: MRPL15*, yellow)*, C2_07680W* (*S.c.: MRPL7*, purple)*, CR_01370C* (*S.c.: MRPS28*, blue)*, and CR_04140W* (*S.c.: NAM9*, green).

**Figure S6:**
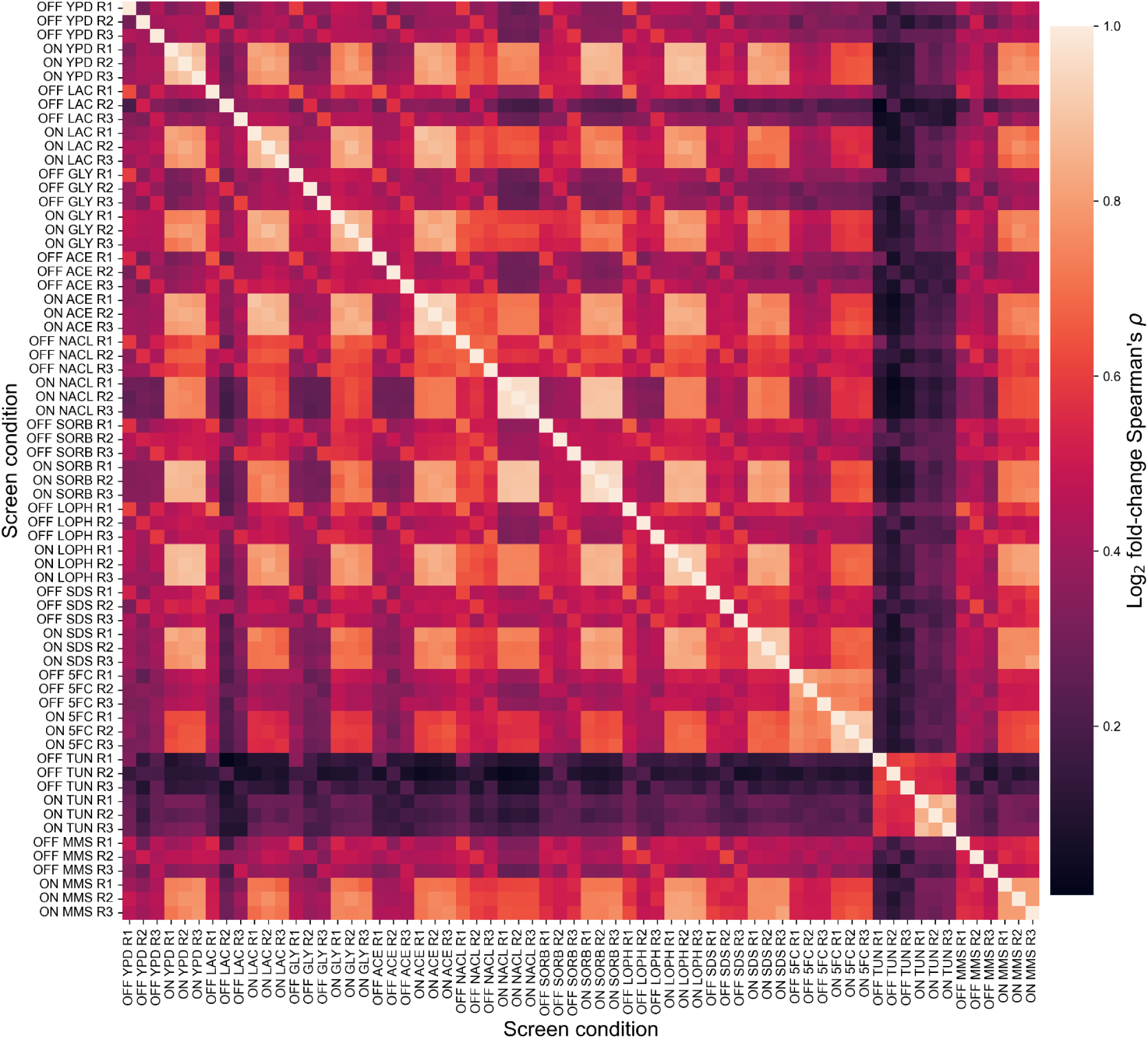
Inter-replicate and inter-condition correlation in the multi-condition screen. Spearman’s rank correlation matrix across all samples (n=469 sgRNAs per comparison) for CRISPRi-OFF and CRISPRi-ON samples.

**Figure S7:**
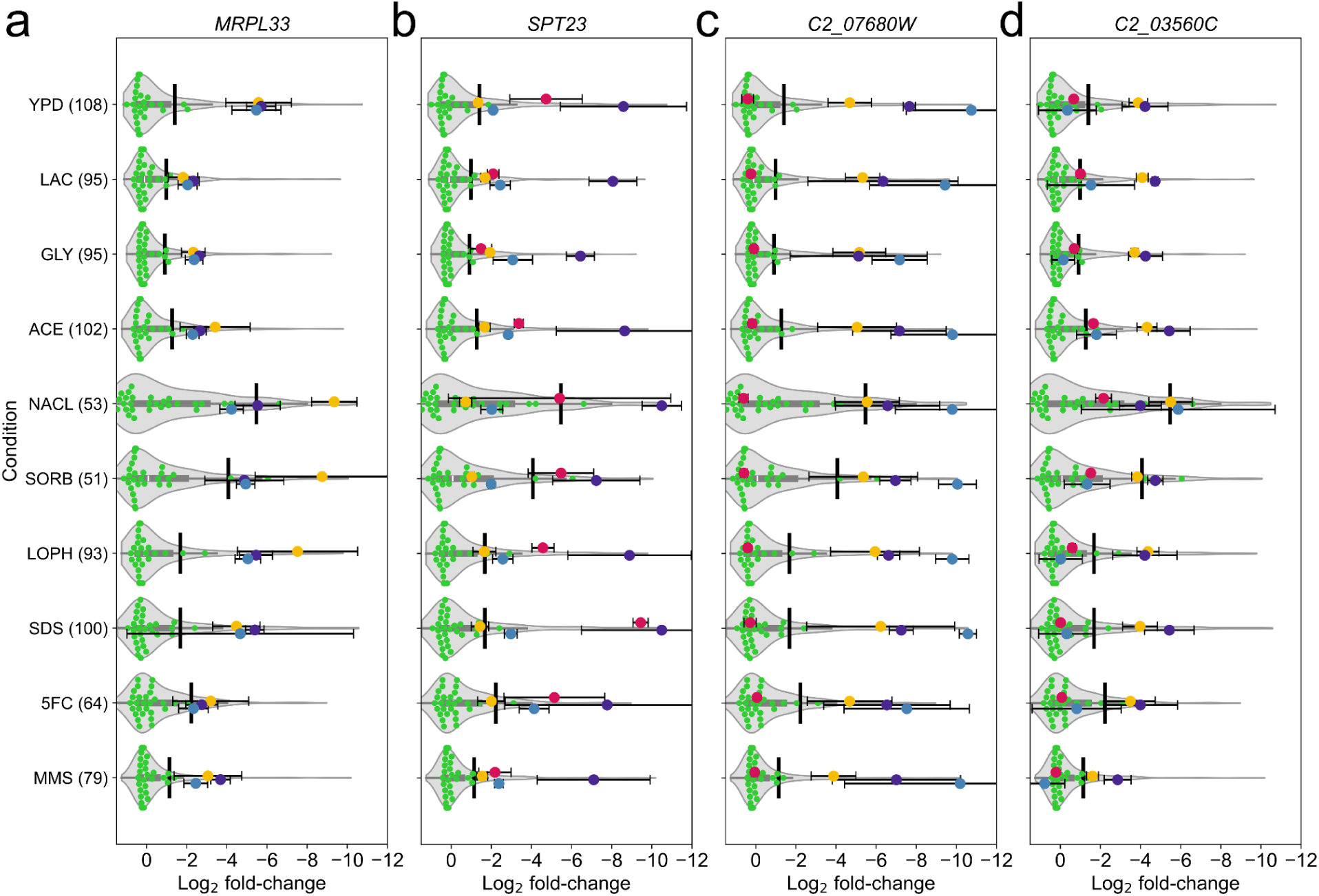
Genes with constant repression sensitivity phenotypes. For each gene, the overall distribution of Log_2_FC for library sgRNAs for each condition is shown as a violin plot, with the randomized sgRNAs shown individually in green. The black lines indicate the significance threshold for each condition. Error bars show the 95% confidence interval around the mean Log_2_FC. **a)** *MRPL33* **b)** *SPT23* **c)** *C2_07680W* **d)** *C2_03560C*.

**Figure S8:**
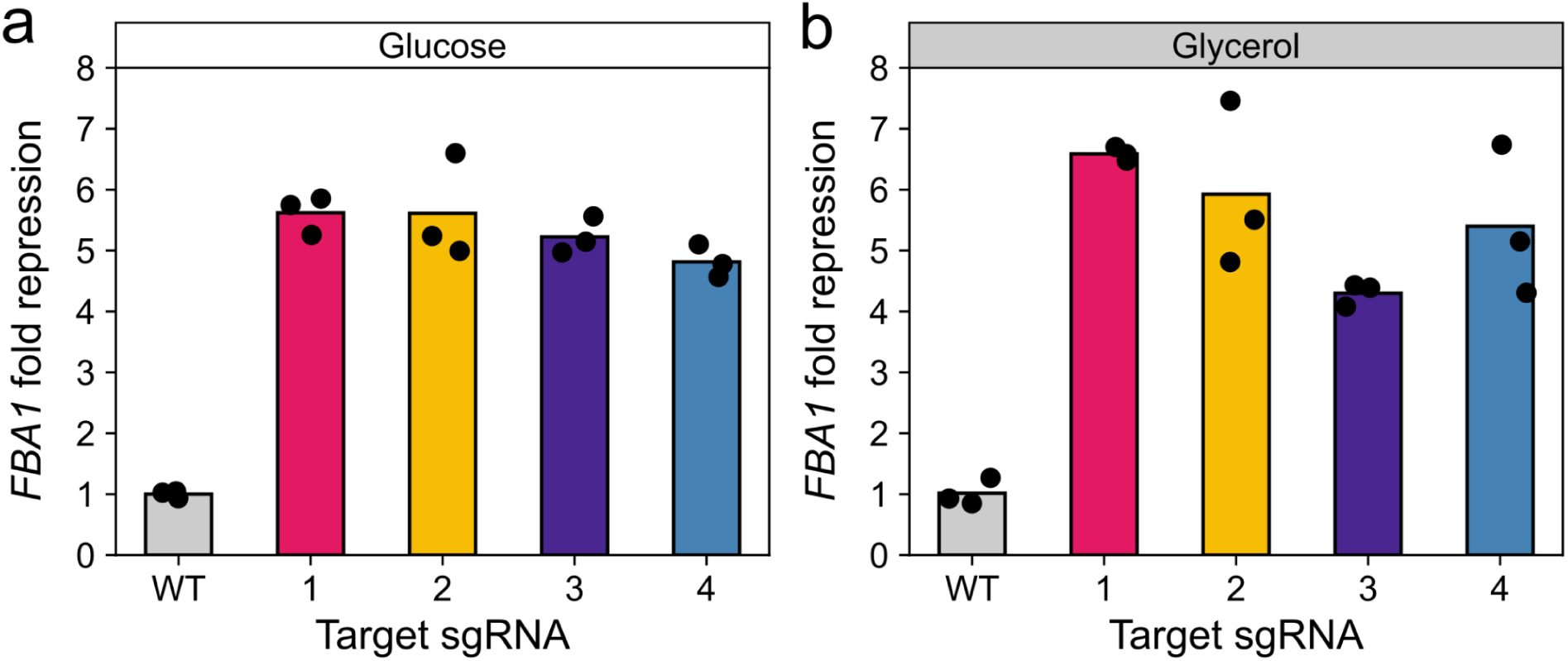
CRISPRi repression of *FBA1* is comparable in glucose and glycerol conditions. CRISPRi repression of *FBA1* in media with **a)** glucose or **b)** glycerol as the carbon source.

**Figure S9:**
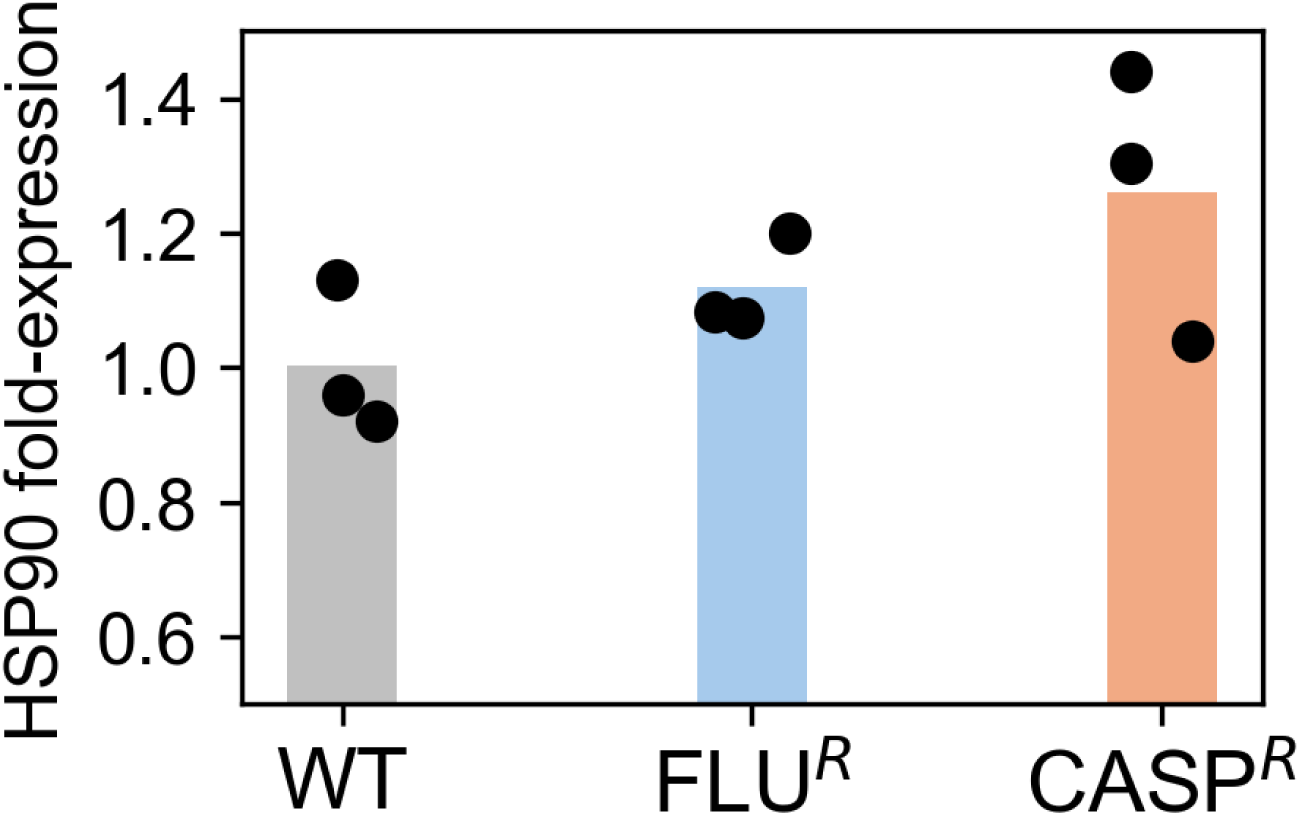
Baseline *HSP90* expression levels in the three strains of interest. Each dot represents a different biological replicate. Difference in expression compared with the wild-type was tested using a two-sided Welch’s t-test: WT vs FLU^R^: p=0.22, WT vs CASP^R^: p=0.15.

**Figure S10:**
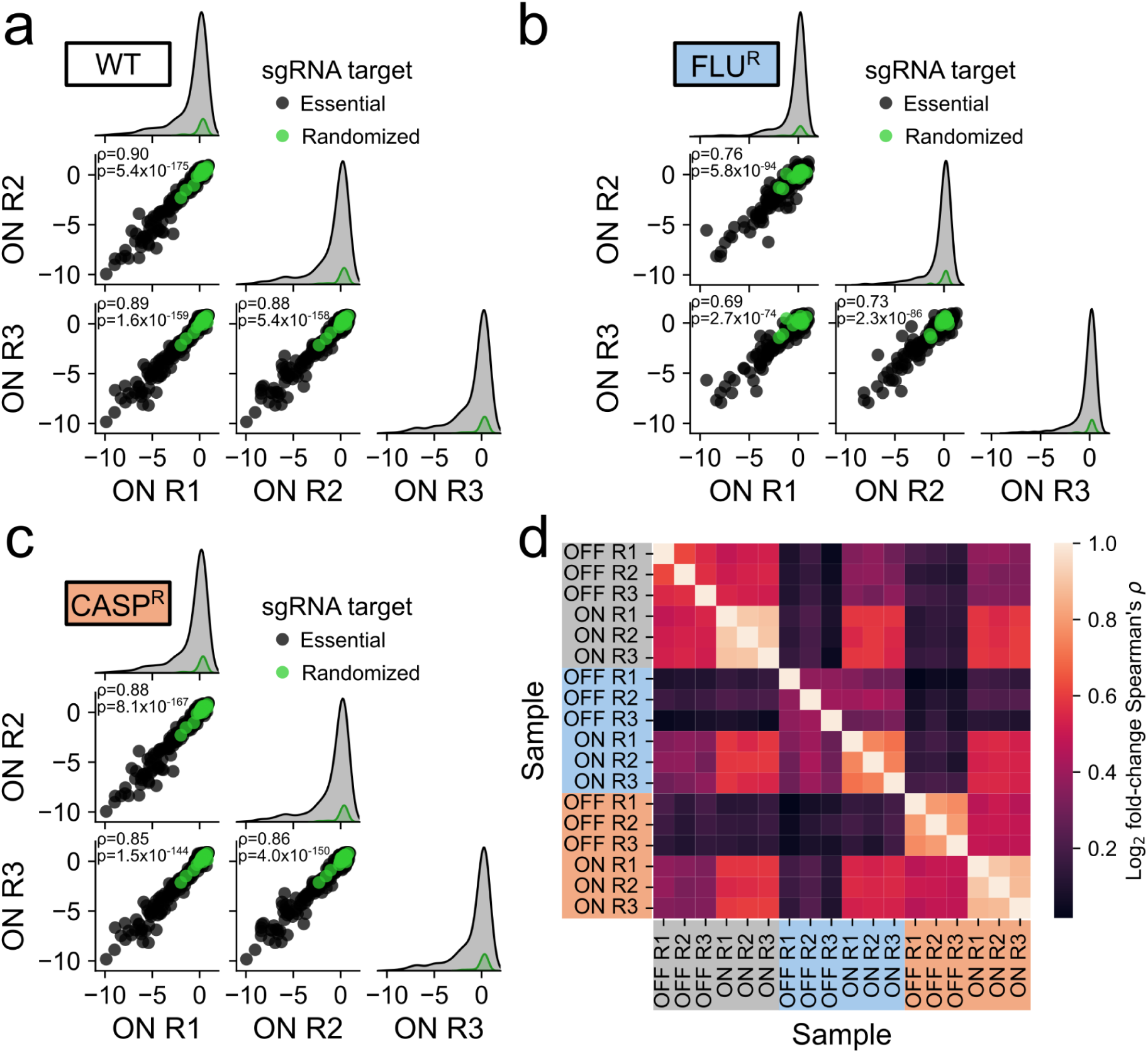
Pooled CRISPRi screening is reproducible in different genetic backgrounds. **a)** Inter-replicate correlation for CRISPRi-ON samples of the wild-type strain library (R1 vs R2: n=475, R1 vs R3: n=473, R2 vs R3: n=474). **b)** Inter-replicate correlation for CRISPRi-ON samples of the FLU^R^ strain library (R1 vs R2: n=526, R1 vs R3: n=515, R2 vs R3: n=515). **c)** Inter-replicate correlation for CRISPRi-ON samples of the CASP^R^ strain library (R1 vs R2: n=521, R1 vs R3: n=519, R2 vs R3: n=519). d) Spearman’s rank correlation matrix across all backgrounds (n=436 sgRNAs per comparison) for CRISPRi-OFF and CRISPRi-ON samples.

**Figure S11:**
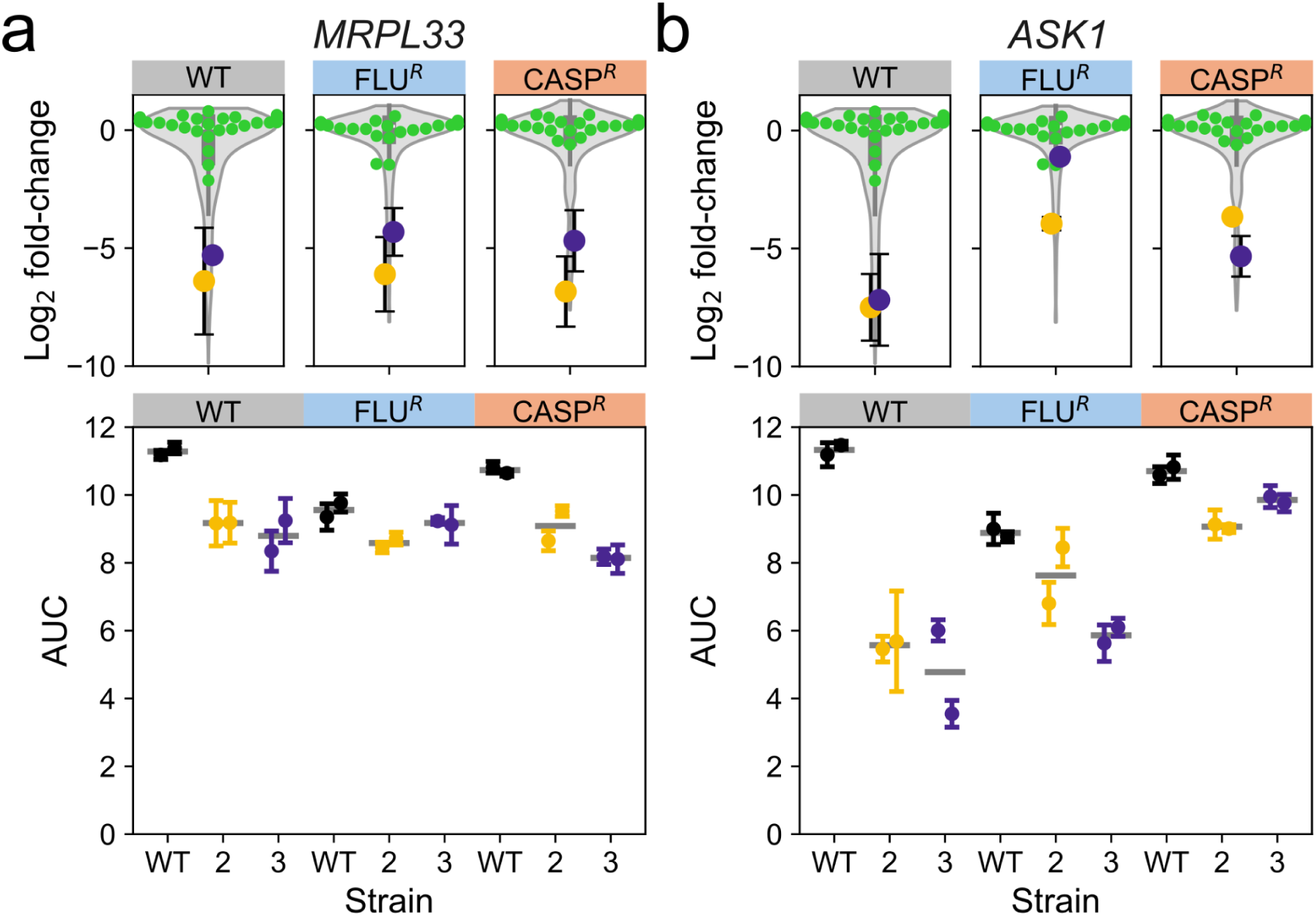
Fitness effects of depleted sgRNAs can be validated across backgrounds. Fitness effects of sgRNAs in the pooled assay (top) and in small-scale growth curve assays (bottom). The overall distribution of Log_2_FC for library sgRNAs is shown as a violin plot, and the randomized sgRNAs are shown individually in green. **a)** Effects for *MRPL33-2* (yellow) and *MRPL33-3* (purple) **b)** Effects for *ASK1-2* (yellow) and *ASK1-3* (purple).

**Figure S12:**
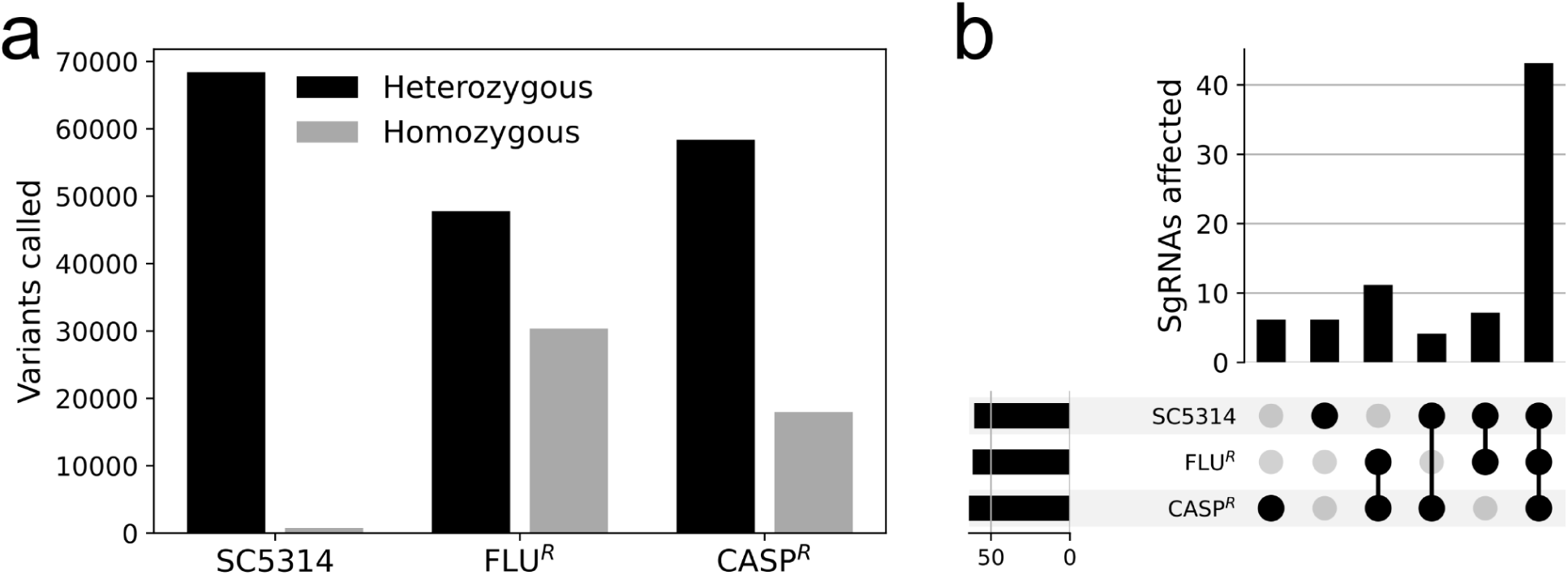
Few sgRNAs are affected by heterozygous or homozygous variants. **a)** Single nucleotide polymorphisms (SNPs) detected by whole genome sequencing for the three strains when compared to the SC5314 reference genome. **b)** Overlap between sgRNAs with target binding regions affected by SNPs. The lower left histogram shows the total number of sgRNAs affected for each strain.

**Figure S13:**
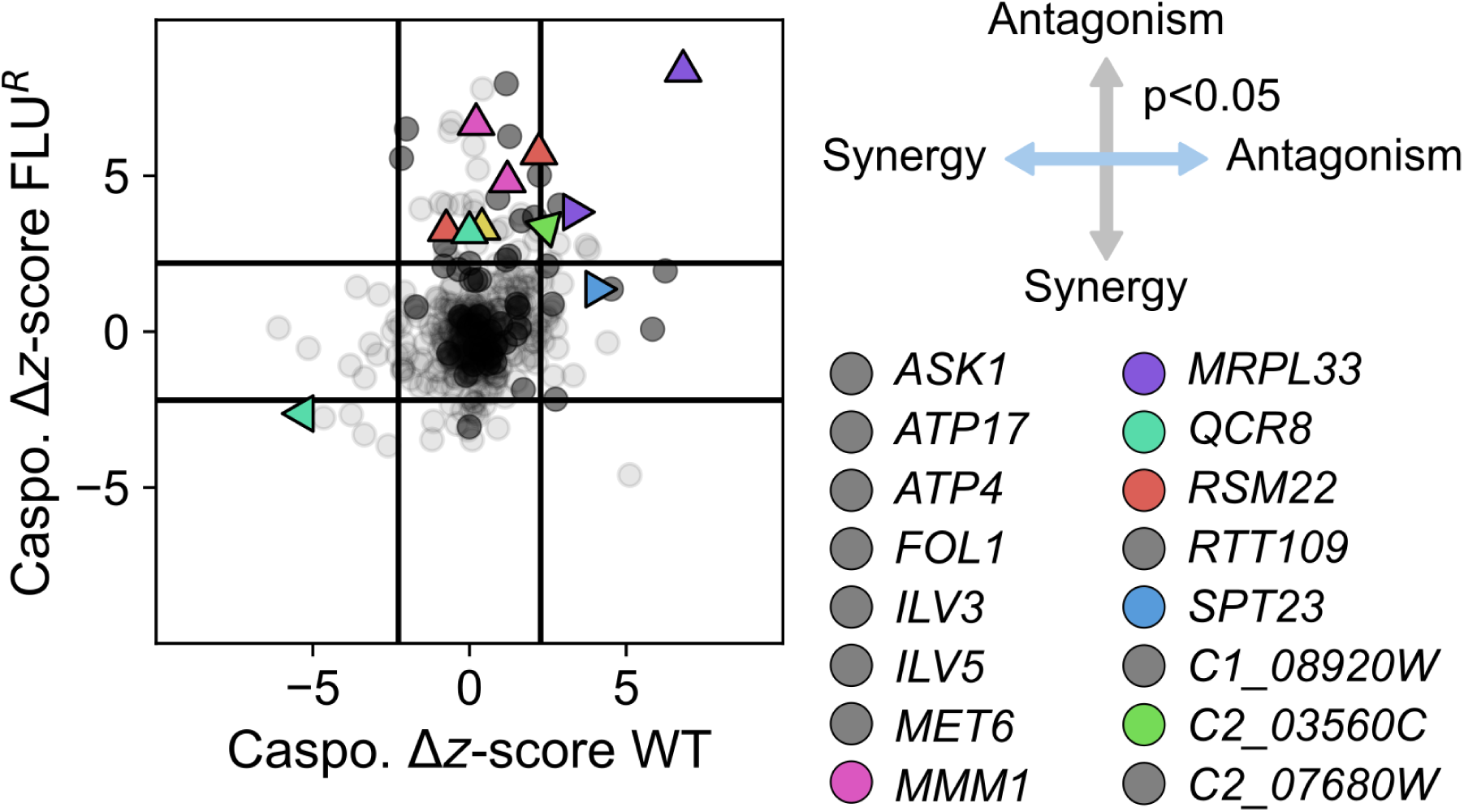
CRISPRi antagonism or synergy with caspofungin in the FLU^R^ genetic background. Interaction between CRISPRi repression and Caspofungin treatment. Only genes with at least one sgRNA showing antagonism or synergy of sufficient magnitude and statistical significance (Benjamini-Hochberg corrected p-value <0.05) are shown as triangles and colored. The direction of the triangle indicates the nature of the interaction, as indicated by the compass: left: synergy in WT (*QCR8*); right: antagonism in WT (*MRPL33*, *SPT23*); down: synergy in CASP^R^ (none); up: antagonism in CASP^R^ (*MMM1*, *MRPL33*, *QCR8*, *RSM22*, *SPT23*); upper right: antagonism in both strains (*C2_035560C*). The black lines show the thresholds for the magnitude of z-score change (see methods). Genes and sgRNAs not showing synergy or antagonism are shown in grey.

## Supplementary Note 1: Outline of procedures to build *Candida albicans* CRISPRi pooled libraries

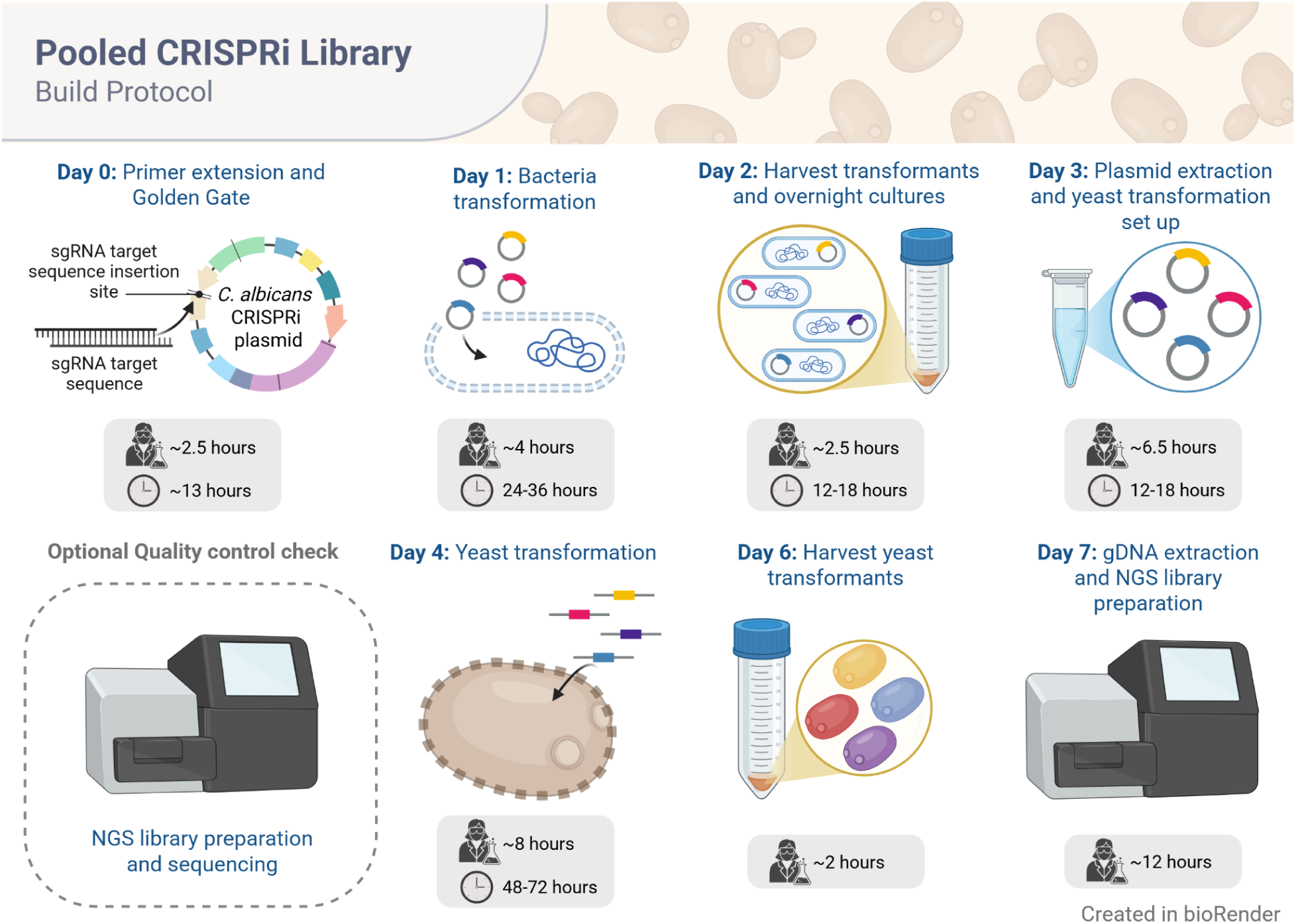

### Before starting: Design a custom sgRNA library

1. Design sgRNAs according to experimental goals (see methods: *sgRNA library design*).
2. Order pooled oligonucleotides libraries from suppliers (In this work, we used IDT or Twist Bioscience oligonucleotide pools).

- **TIMING** 3-5 days to design; ∼4-7 weeks for synthesis and delivery (dependent on pool size and supplier).

### Day 0: sgRNA cloning

- Grow bacteria containing the plasmid backbone and extract the plasmid DNA.
- Convert ssDNA oligonucleotide pool to dsDNA (see methods: *Primer extension*).
- Clone pooled oligonucleotides into the CRISPRi plasmid backbone (see methods: *Golden Gate cloning*).

- **TIMING** *Recommended start time 2:00 pm.* Primer extension protocol ∼1.5 hours; Golden Gate protocol ∼1 hour; Thermocycler for ∼13 hours.

### Day 1: Bacteria transformation

- Transform the cloned plasmid pool into *E. coli* DH5α cells (see methods: *Bacteria Transformation*).
- During incubation steps in the bacteria transformation protocol, prepare the LB semi-solid agar for growing transformants.

- **TIMING** *Recommended start time 9:00 am.* Bacteria transformation protocol ∼4 hours; Semi-solid agar preparation ∼1.5 hours; Incubation period 24-36 hours or until colonies become visible.

### Day 2: Harvest Bacteria pooled library

- Harvest bacteria colonies (see methods: *Bacteria Transformation*).
- Prepare an overnight culture for plasmid extraction (see methods: *Plasmid pool extraction*).

- **TIMING** *Recommended start time 3:00 pm.* Harvesting bacteria cells ∼2 hours; Starting overnight culture ∼30 min; Incubation period ∼12-18 hours.

### Day 3: Extract pooled plasmid Library and preparation for Fungal transformation

- Perform plasmid DNA extraction to obtain a large quantity of plasmid (see methods: *Plasmid pool extraction*).
- Linearize pooled plasmids by restriction digest (see methods: *Candida albicans transformation*).
- Prepare an overnight culture of the selected strain background (see methods: *Candida albicans transformation*).

- **TIMING** *Recommended start time 9:00 am.* Plasmid extraction protocol ∼5 hours; Overnight culture set-up, and restriction enzyme digest ∼1.5 hours; Incubation period ∼12-18 hours.

### Optional (Highly recommended) Sequencing of the pooled plasmid library

- After extracting the plasmid library, you can perform high-throughput sequencing to determine the composition of your plasmid library and validate adequate representation before moving on to fungal transformation steps (see methods: *NGS library preparation and sequencing*).

- **TIMING** *Recommended start time 9:00 am.* Library preparation for sequencing ∼8 hours; Sequencing turnaround time dependent on the provider.

### Day 4: *Candida albicans* transformation

- Transform *C. albicans* with the pooled CRISPRi plasmid library (see methods: *Candida albicans transformation*).
- Prepare the YPD semi-solid agar during incubation steps.

- **TIMING** *Recommended start time 8:00 am.* Fungal transformation protocol ∼8 hours; Semi-solid agar protocol ∼1.5 hours; Incubation period 48-72 hours or until colonies become visible.

### Day 6: Harvest fungal pooled library

- Harvest fungal library (see methods: Candida albicans transformation).
- Prepare frozen aliquots of the fungal library for storage and downstream assays.

- **TIMING** *Recommended start time 4:00 pm.* Harvesting fungal cells ∼2 hours.

### Day 7-8: Sequencing of the pooled plasmid library

- Extract Genomic DNA from a sample of your pooled fungal library (see methods: *Genomic DNA extraction*).
- Prepare samples for sequencing (see methods: *NGS library preparation and sequencing*).

- **TIMING** *Recommended start time 9:00 am.* Genomic extractions ∼4 hours. Library preparation for sequencing ∼8 hours.

